# Mitochondrial, cell cycle control and neuritogenesis alterations in an iPSC-based neurodevelopmental model for schizophrenia

**DOI:** 10.1101/2020.09.04.282046

**Authors:** Giuliana S. Zuccoli, Juliana M. Nascimento, Ana Campos Codo, Pedro M. Moraes-Vieira, Stevens K. Rehen, Daniel Martins-de-Souza

## Abstract

Schizophrenia is a severe psychiatric disorder of neurodevelopmental origin that affects around 1% of the world’s population. Proteomic studies and other approaches have provided evidence of compromised cellular processes in the disorder, including mitochondrial function. Most of the studies so far have been conducted on postmortem brain tissue from patients and do not allow the evaluation of the neurodevelopmental aspect of the disorder. To circumvent that, we studied the mitochondrial and nuclear proteomes of neural stem cells (NSCs) and neurons derived from induced pluripotent stem cells (iPSCs) from schizophrenia patients versus healthy controls. Our results revealed differentially regulated proteins in pathways related to mitochondrial function, oxidative phosphorylation, cell cycle control, DNA repair, and neuritogenesis. Moreover, metabolic analysis of NSCs revealed alterations in mitochondrial function in schizophrenia-derived cells. Hence, this study shows that changes in important cellular processes are present during neurodevelopment and could be involved with the establishment of schizophrenia, as well as the phenotypic traits observed in adult patients.

## Introduction

Schizophrenia is a complex syndrome expressed by the combination of psychotic symptoms, such as hallucinations and delusions, with cognitive and motivational dysfunctions (Kahn et al., 2015). The disorder affects approximately 1% of the world’s population and has a neurodevelopmental origin, which indicates that genetic and environmental factors promote early insults accompanied by a latent period throughout neurodevelopment, leading to the appearance of psychosis on late adolescence (Insel, 2010).

Proteomic studies conducted on postmortem brain tissue have provided insights on cellular processes that may be altered in the disorder, such as inflammation, glucose handling, mitochondrial function, and response to reactive oxygen species (ROS) (Martins-de-Souza et al., 2011; Nascimento and Martins-de-Souza, 2015; Zuccoli et al., 2017). Organelle proteomics is a valuable tool and is based on the enrichment of a specific organelle in a given sample prior to mass spectrometry (MS) analysis, which allows better coverage of the subcellular proteome, enabling the identification of less abundant proteins (Taylor et al., 2003).

Most of the molecular studies so far have been conducted on postmortem brain tissue from schizophrenia patients, which makes it impossible to investigate the disorder in living cells and during the different developmental stages of the disorder (Pedrosa et al., 2011). The use of induced pluripotent stem cells (iPSCs) (Takahashi et al., 2007; Takahashi and Yamanaka, 2006) derived from patients with schizophrenia has provided the possibility to study different neural cell types as well as the neurodevelopmental aspect of the disorder (Marchetto et al., 2011). Neural stem cells (NSCs) derived from those iPSCs allow studying the process of neurons and radial glia giving rise to more defined neuronal and glial populations (Marchetto et al., 2010). Studies performed on NSCs and neural progenitor cells (NPCs) derived from iPSCs from schizophrenia patients showed increased oxygen consumption, decreased angiogenic factors, and synaptic alterations (Brennand et al., 2015, 2011a; Casas et al., 2018; Paulsen et al., 2014, 2012). These characteristics indicate that this model can help recapitulate the phenotypic traits of schizophrenia, but the specific proteins and pathways involved have not yet been elucidated.

Hence, in this work, we proposed to study the mitochondrial and nuclear proteome of NSCs and neurons derived from iPSCs from patients with schizophrenia, as well as the whole proteome of neurons. Mitochondria provide most of the energy for the cell and are involved in ATP production, lipid metabolism, Ca^2+^ buffering and, ROS production. In stem cells, proper mitochondrial function is important for neurogenesis and neural stem cell fate decisions (Khacho and Slack, 2018). It has been proposed that mitochondrial dysfunction is central to the development of schizophrenia (Hroudová and Fišar, 2011; Rezin et al., 2009). For that reason, we aimed at identifying the proteins and pathways altered in the disorder and provide further information regarding the possible alterations related to energy metabolism and its consequences to the neurodevelopmental course of schizophrenia.

## Experimental Procedures

### Generation of iPSCs from schizophrenia patients and controls

Schizophrenia cell lines used in this study were obtained from three subjects diagnosed in the schizophrenia spectrum, GM23760B (SCZ 1), GM23761B (SCZ 2), and EZQ4 (clone 1) (SCZ 3). Patient 1 (GM23760B) and patient 2 (GM23761B) are siblings (Brennand et al., 2011b) (available at Coriell). Three control cell lines were used GM23279A (available at Coriell) (CTRL 1), CF1 (clone 10) (CTRL 2) and CF2 (clone 2) (CTRL 3). Cell lines EZQ4, CF1, and CF2 were reprogrammed at the D’Or Institute for Research and Education (Sochacki et al., 2016). Cells were cultured in mTeSR1 media (Stemcell Technologies, Vancouver, Canada) or E8 (Thermo Fisher Scientific, Carlsbad, CA, USA) on Matrigel (BD Biosciences, San Jose, CA, USA)-coated surface. Colonies were manually passaged every five-seven days and maintained at 37°C in humidified air with 5% CO_2_. Further information regarding the reprogramming of fibroblasts into iPSCs are described in Casas et al., 2018 (Casas et al., 2018).

### Neural differentiation

iPSCs lines derived from schizophrenia patients and controls were adapted to E8 medium (Thermo Fisher Scientific, Carlsbad, CA, USA) for a minimum of 4 passages and the cells were then split. After 24 h of splitting the cells, we maintained them in Pluripotent Stem Cells (PSC) Neural Induction Medium (Thermo Fisher Scientific, Carlsbad, CA, USA), composed by Neurobasal medium and PSC supplement, according to the manufacturer’s protocol. After seven days, changing the medium every other day, cells were then maintained in Neural Induction Medium (NEM, Advanced DMEM/F12, and Neurobasal medium (1:1) with Neural Induction Supplement; Thermo Fisher Scientific, Carlsbad, CA, USA).

### NSCs differentiation into neurons

NSCs were plated on poly-ornithine/laminin-coated plates and maintained with NEM medium at 37°C in humidified air with 5% CO_2_. Upon reaching 40% confluency, the medium was changed to the neuronal differentiation medium, which consists of DMEM/F12 and Neurobasal medium (1:1) with 1% B27 supplement (Thermo Fisher Scientific, Carlsbad, CA, USA). The medium was changed every five days for 30 days and consisted of the removal of half of the content and addition of this same volume of fresh medium. Factors secreted by the differentiating cells are important for a successful differentiation.

### Subcellular fractionation

Mitochondrial and nuclear fractions were obtained from NSCs and neurons according to an adaptation of the protocol described by Clayton and Shadel (Clayton and Shadel, 2014). Approximately 10^7^ cells from each group were homogenized on ice in a buffer containing 210 mM Mannitol (Sigma-Aldrich, St. Louis, MO, USA), 70 mM Sucrose (Sigma-Aldrich), 5 mM Tris-HCl (Sigma-Aldrich), 1 mM EDTA (Sigma-Aldrich) and one tablet of protease cocktail inhibitor (Roche Diagnostics, Indianapolis, IN, USA) per 25 ml of buffer. The homogenized samples were then centrifuged at 2000xg for 5 minutes. The resulting pellets consisted of the nuclear fraction. The supernatant was then centrifuged at 13000xg for 20 minutes and the resulting pellets were the mitochondrial fraction.

### In-gel digestion

Mitochondrial, nuclear, and whole-cell pellets were subjected to a brief sodium dodecylsulfate polyacrylamide gel electrophoresis (SDS-PAGE) to enhance the protein separation in each sample and optimize the digestion into peptides. Gel sections containing the proteins were subjected to reduction, alkylation, and overnight digestion with trypsin as described by Shevchenko et al. (Shevchenko et al., 2007). The resulting peptide mixtures were removed from the gel by addition of acetonitrile and the solution with peptides were dried and posteriorly resuspended in ammonium formate pH 10 to posterior LC-MS/MS analysis.

### LC-MS/MS

Proteomic analyses were performed in a bidimensional nanoUPLC tandem nanoESI-HDMSE platform by multiplexed data-independent acquisitions (DIA) experiments. The peptides (1 μg) were injected into a 2D-RP/RPAcquity UPLC M-Class System (Waters Corporation, Milford, MA) coupled to a Synapt G2-Si mass spectrometer (Waters Corporation, Milford, MA). The samples were fractionated in first dimension chromatography with an XBridge Peptide BEH C18 NanoEase Column (Waters Corporation, Milford, MA). Peptide elutions were carried to second dimension separation in an ACQUITY UPLC HSS T3 nanoACQUITY Column (Waters Corporation, Milford, MA). They were achieved by using an acetonitrile gradient from 7% to 40% (v/v) for 95 min at a flow rate of 500 nL/min directly into a Synapt G2-Si. MS/MS analyses were performed by nano-electrospray ionization in positive ion mode nanoESI (+) and a NanoLock Spray (Waters, Manchester, UK) ionization source.

### Database search and protein identification

Spectra processing and database searching conditions were performed on the Progenesis^®^ QI version 3.1 software package with Apex3D, peptide 3D, and ion accounting informatics (Waters). Proteins were identified and quantified by searching on the Uniprot human proteomic database, version 2017/10, with the following parameters for peptide identification: 1) trypsin digestion with up to one missed cleavage; 2) variable modifications by oxidation (M) and fixed modification by carbamidomethyl (C); 3) false discovery rate (FDR) less than 1%; and 4) peptides/proteins (N) equal to 3 in the quantification of proteins by the method of relative quantification with HI\N. Identifications that did not meet these criteria weren’t considered.

### Systems biology analysis in silico

Biological processes and cellular compartments from each protein were searched in the Human Protein Reference Database (https://hprd.org/). Gene ontology analysis of biological processes related to the differentially regulated proteins was performed using the David database (http://david.ncifcrf.gov). Canonical pathways associated with differentially expressed proteins were identified by Ingenuity Pathway Analysis (IPA, Ingenuity Systems, Qiagen, Redwood, CA, USA; www.ingenuity.com). This program is based on an algorithm that uses curated connectivity information from the literature to determine the network of interactions among the differentially expressed proteins and canonical pathways in which they are involved (Calvano et al., 2005).

### Immunofluorescence

NSCs were plated on geltrex-coated glass coverslips (13 mm) at a density of 4×10^6^ cells/coverslip. NSCs were grown until 80% confluency was reached. For neurons, NSCs were differentiated on coverslip following the previously described differentiation protocol. NSCs and 30-day neurons were then fixed with 4% paraformaldehyde (PFA) for 10 minutes and incubated for 10 minutes in the permeabilization solution (0.1% Triton-X-100 in PBS). Followed by a blocking solution (1% bovine serum albumin in PBS) for 30 minutes. NSCs were incubated with the primary antibody anti-nestin (1:200; Cell signaling, #33475) and neurons were incubated with the primary antibody anti-β3-tubulin (1:200; Cell signaling, #4466). Primary antibodies were diluted in blocking solution and incubated overnight in a humid chamber at 4°C. On the next day, coverslips were incubated with the secondary antibody diluted in blocking solution (1:400; Alexa Fluor^®^ 488 Conjugate, #4408) and for the nuclear stain, DAPI (Cell signaling, #4083) was used on the final 10 minutes of incubation. Slides were then mounted with Neo-Mount (Sigma-Aldrich) and viewed on Cytation™ 5 Imaging Multi-mode Reader (BioTek).

### NSCs metabolic analysis

NSCs (SCZ and CTRL) were plated at a density of 9×10^4^ per well on Seahorse Flux™ 24 plates and incubated at 37°C in 5% CO_2_. 24h later, for the Mito Stress Test, cell medium was changed to Seahorse Base medium (supplemented with 1mM pyruvate, 2mM glutamine, and 10mM glucose) and the plate was incubated for one hour at 37°C without CO_2_. For the experiment, OCR (Oxygen Consumption Rate) was measured under basal conditions and after the additions of oligomycin (0.5μM), FCCP (1.5μM), and rotenone/antimycin A (1μM). For the Glycolysis Stress Test, the cell medium was changed to Seahorse base medium (supplemented with 2mM glutamine) and the plate was incubated for one hour at 37°C without CO_2_. For the experiment, ECAR (Extracellular Acidification Rate) was measured under basal conditions and after the additions of glucose (10mM), oligomycin (0.5μM) 2-Deoxi-d-glucose (5mM). Measurements for each well were normalized by protein concentration.

### Reactive Oxygen Species

Reactive oxygen species levels were estimated using the probe CM-H2DCFDA (Molecular probes, Invitrogen). NSCs (SCZ and CTRL) were plated at a density of 6×10^4^ cells per well in a 96 well plate on the day before the experiment. Cells were incubated with 10μM CM-H2DCFDA probe for 30 min at 37°C and then washed twice with PBS. Fluorescence (485/535) was measured immediately using Cytation™ 5 Imaging Multi-mode Reader (BioTek). Measurements for each well were normalized by protein concentration (Oparka et al., 2016).

## Results and discussion

### Mitochondrial proteome and metabolic analysis of NSCs derived from SCZ iPSC

NSC’s (Figure 1, A and B) mitochondrial proteome analysis revealed differentially regulated proteins (Sup. Table 1) related to oxidative phosphorylation, calcium homeostasis, and *cristae* formation. With mitochondrial dysfunction canonical pathway significantly dysregulated in NSC (p=5.04E-05).

**Figure 1:**
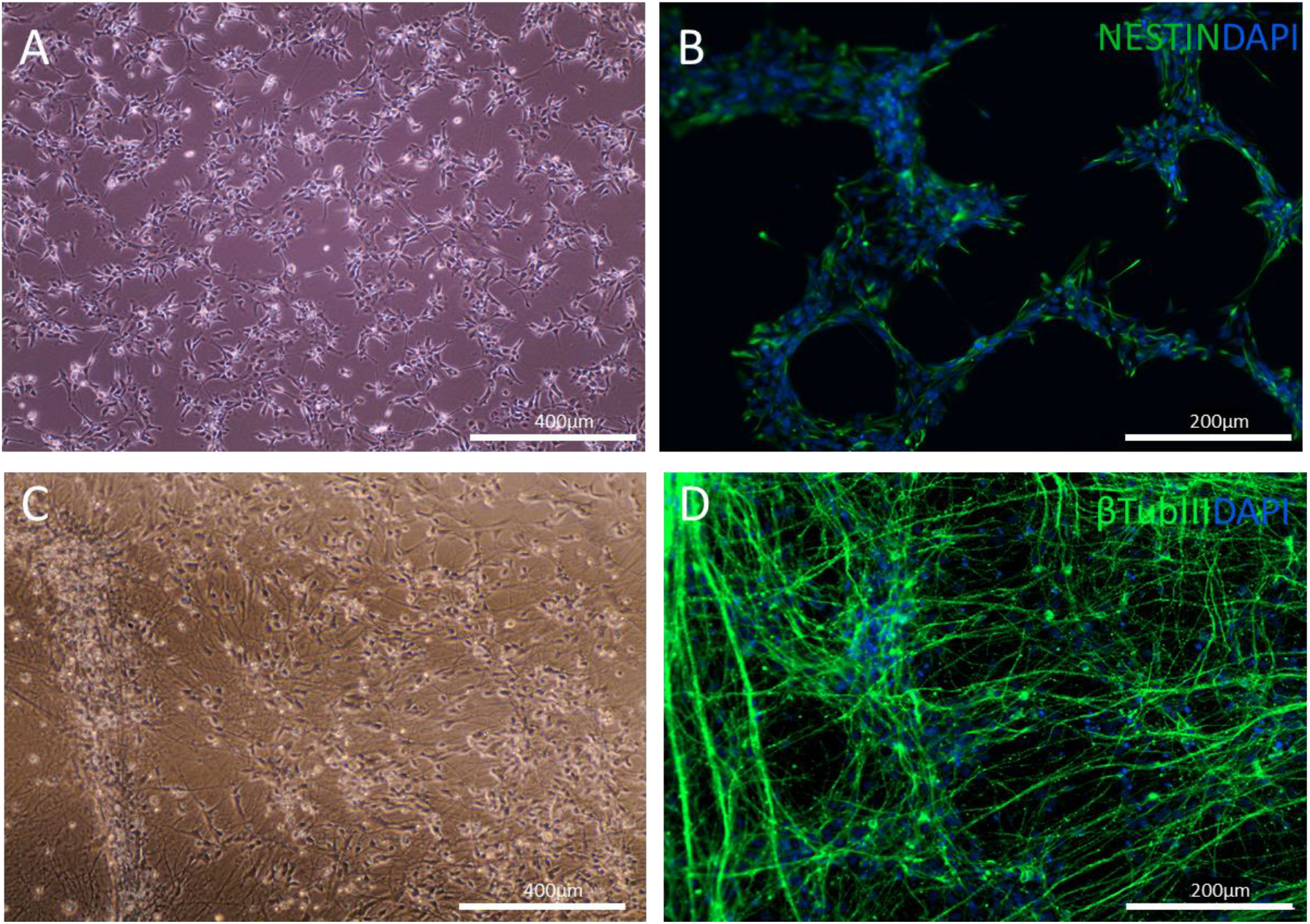
NSCs and Neurons derived from iPSC. Representative image of iPSC-derived NSCs (A) and immunostaining for the neural stem cell marker Nestin (B). Representative image of iPSC-derived neurons (C) and immunostaining for the neuronal marker β-tubulin III (D). DAPI was used as nuclear marker.

Hence, to evaluate the bioenergetic profile of those cells, we assessed the mitochondrial and glycolytic function of NSCs. Where we have found reduced levels of non-mitochondrial oxygen consumption and elevated levels of basal respiration, maximal respiration, ATP production, and spare respiratory capacity when comparing schizophrenia and control samples (Figure 2). Thus, revealing alterations in key parameters of mitochondrial function but not in parameters related to glycolytic metabolism. In addition, by assessing intracellular ROS formation with CM-H2DCFDA probe staining, our data suggests a tendency of higher levels of ROS in SCZ-NSCs when compared to CTRL-NSCs (Figure 3), which corroborate the proteomic alterations found and previous results obtained with patient’s NPCs (Paulsen et al., 2012).

**Figure 2:**
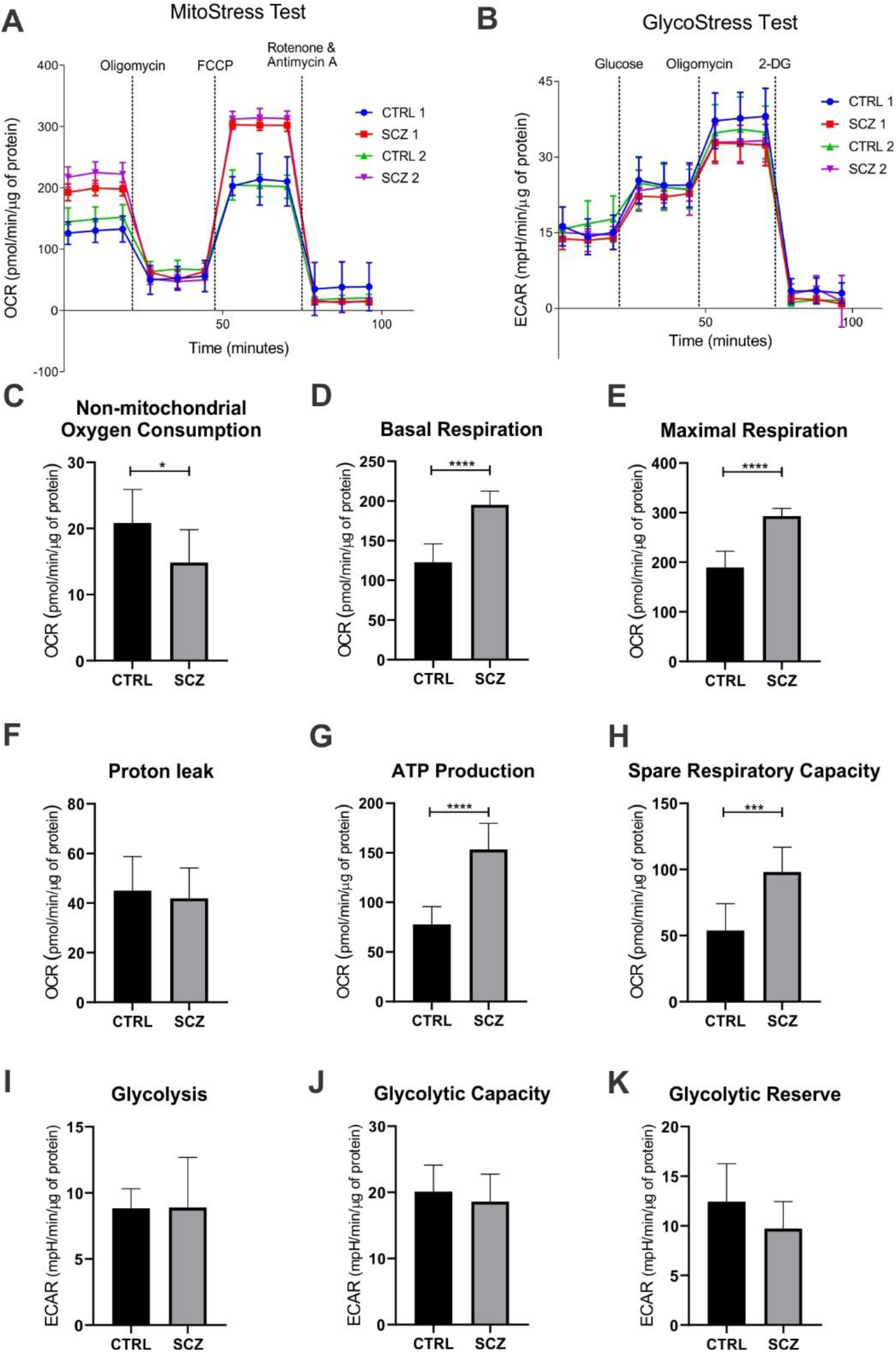
Mitochondrial and glycolytic metabolic analysis of NSCs from schizophrenia patients and controls. (A) Oxygen consumption rate and (B) Extracellular acidification rate measured throughout the experiment. From which were calculated the (C) Non-mitochondrial oxygen consumption, (D) Basal respiration, (E) Maximal respiration, (F) Proton leak, (G) ATP production, (H) Spare respiratory capacity, (I) Glycolysis, (J) Glycolytic capacity, and (K) Glycolytic reserve. Data shown are means±SD. n=5 Welch’s ANOVA test: *p < 0.05, ***p < 0.001, **** p < 0.0001.

**Figure 3:**
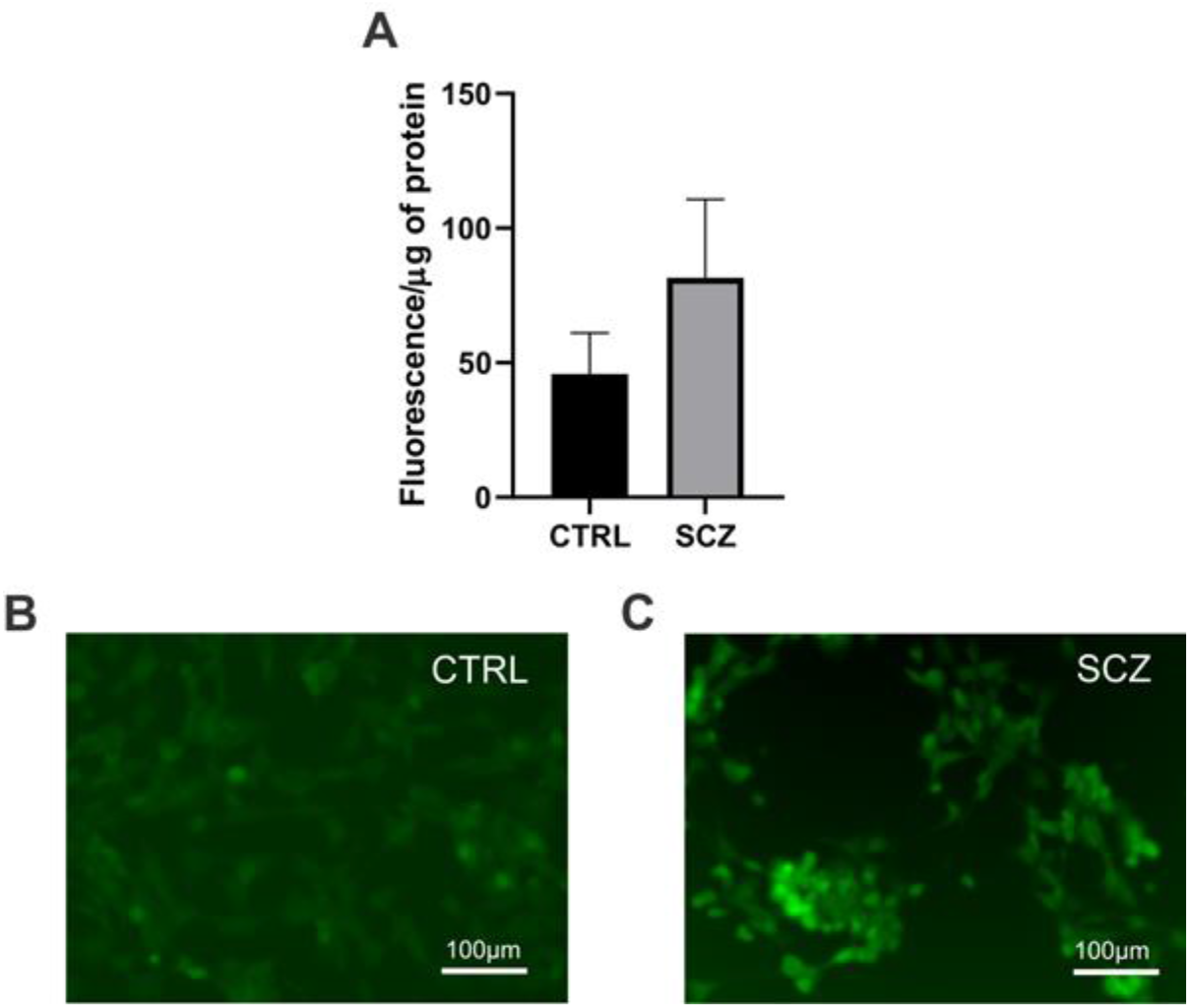
Intracellular ROS formation. (A) ROS levels detected in NSCs normalized by protein concentration. n=5 Welch’s ANOVA test was performed (p=0.0508). Immunofluorescence staining of CTRL (B) and SCZ (C) cells.

NSCs display a highly glycolytic metabolism and have a low demand for oxygen metabolism (Candelario et al., 2013). Given this low oxidative need, proper oxygen tension, and redox state are modulators of neural stem cell metabolism, fate commitment, and survival (Panchision, 2009; Pistollato et al., 2007; Studer et al., 2000). Therefore, alterations in mitochondrial metabolism found in this study point to cellular disturbances, since despite not being essential for ATP production in NSCs, mitochondrial function is required for stem cell self-renewal (Khacho et al., 2016; Khacho and Slack, 2017; Steib et al., 2014). Also, regulation of oxidative metabolism and ROS levels determines the maintenance of stem cell trait, whereas increased ROS levels via enhanced oxidative phosphorylation determine stem cell differentiation (Khacho et al., 2016). Our results showed altered regulation of proteins related to cristae formation in schizophrenia cells, and changes in mitochondrial morphology in NSCs act as an upstream regulatory mechanism for stem cell fate decisions through modification of ROS signaling (Khacho et al., 2016; Steib et al., 2014). In this way, the NSCs from schizophrenia patients and controls analyzed in this study presented proteomic and functional differences in important mitochondrial processes that are associated with proper development.

### Nuclear proteome of NSCs and neurons derived from SCZ iPSCs

The nuclear NSCs analysis revealed differentially expressed proteins (Sup. Table 1) involved in DNA repair and mRNA splicing, with alterations in the canonical pathways cell cycle control (p=1.18E-06), and DNA double-strand break repair (p= 8.15E-06). The control of cell cycle progression is crucial for the generation of appropriate cell fates during neurodevelopment (Cremisi et al., 2003). Moreover, a study conducted on NSC culture derived from nasal biopsies from schizophrenia patients revealed alterations in genes and proteins related to cell cycle control, hypothesizing that subtle alterations in dynamics and shortening of the cell cycle in NSCs could contribute to the neurodevelopmental aspect of the disorder (Fan et al., 2012). Also, several studies have linked polymorphisms in genes related to DNA repair to the development of schizophrenia (Odemis et al., 2016; Saadat et al., 2008) and deficiencies in DNA repair mechanisms in the nervous system could compromise proper developmental dynamics (Lee and McKinnon, 2007).

In addition, our analysis revealed a potential link between nuclear proteins found altered in NSCs and the involvement of Akt and GSK3 in this neurodevelopmental stage in schizophrenia (Figure 4C). Akt acts as a positive regulator of several signaling pathways such as cell proliferation, growth, survival, and metabolism (Engelman et al., 2006). GSK3 proteins were originally identified as regulatory enzymes in glucose metabolism (Woodgett and Cohen, 1984) but have emerged as regulators of neurodevelopmental processes, including neurogenesis, neuronal migration, and axonal guidance (Hur and Zhou, 2010). Besides, evidence suggests a critical role for Akt/GSK3 in the preservation of mitochondrial integrity, function, and morphology (Ong et al., 2015; Wang et al., 2017). Several studies have provided evidence of the involvement of the Akt/GSK3 signaling pathway in schizophrenia (Emamian, 2012) and our work has linked the proteins involved in this pathway to neurodevelopment in the disorder and potentially to the mitochondrial alterations observed in NSCs.

**Figure 4:**
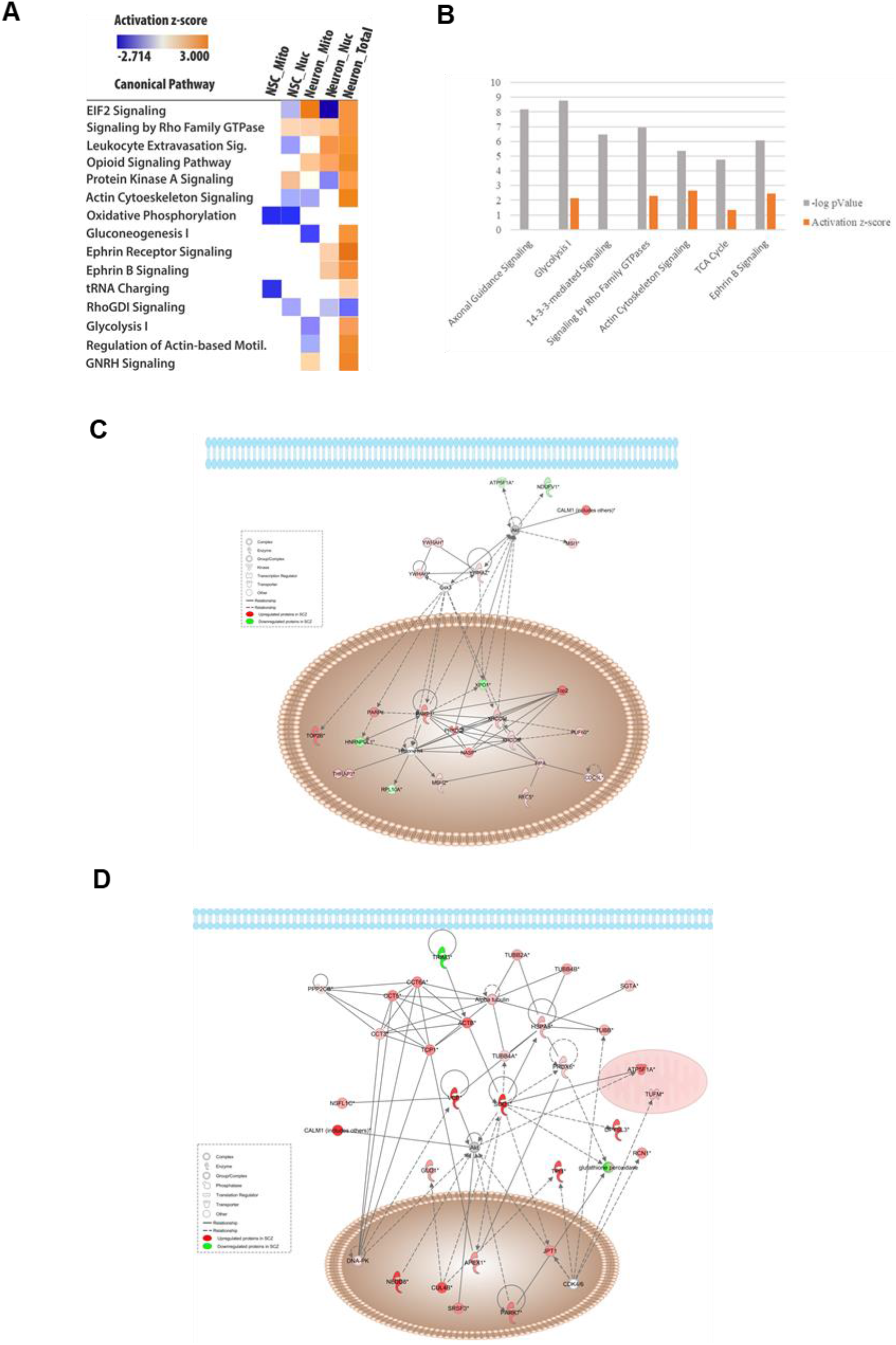
Proteomics analysis regulation and protein interactions of NSC and neurons derived from iPSCs from schizophrenia in comparison to control. (A) Comparison analysis of the levels of activation and inhibition of canonical pathways related to the total, mitochondrial, and nuclear proteome of neurons and the mitochondrial and nuclear proteomes of NSCs from schizophrenia patients in relation to controls. (B) Analysis of the canonical pathways related to the total proteome of neurons derived from iPSCs from schizophrenia patients and their levels of activation. (C) Network analysis of the differentially expressed proteins in the nuclear proteome of NSCs. Colored interactors represent proteins previously found in the proteome. Full and dashed lines depict direct and indirect connections, respectively (D) Network analysis of the differently expressed proteins in neurons derived from schizophrenia patients, showing their interactions and interactors. The network was generated from differentially expressed proteins by IPA. Colored interactors represent proteins previous found in the proteome. Full and dashed lines depict direct and indirect connections, respectively

The nuclear proteome of neurons revealed differentially expressed proteins (Sup. Table 2) related to response do DNA damage, translation, mRNA stability, and cell cycle, with the neuritogenesis (p=5.78E-07) and axonogenesis (p=2.03E-06) canonical pathways significantly altered. The formation of neurites is an essential process in neurodevelopment since it is linked to the formation of axons and dendrites in early neuronal development (Polleux and Snider, 2010). Our findings of the presence of neuritogenesis alterations are consistent with other studies that observed reduced neurite outgrowth in neurons derived from schizophrenia patients (Brennand et al., 2011a; Brennand and Gage, 2011) and in postmortem schizophrenia brain tissue (Jones et al., 2002; Kalus et al., 2000).

### Mitochondrial and total proteome of neurons derived from SCZ iPSCs

Neuron’s mitochondrial proteome analysis revealed differentially regulated proteins (Sup. Table 2) related to calcium transport, mitochondrial location, TCA cycle, and cell redox homeostasis. The canonical pathway TCA cycle was significantly altered in SCZ neurons (p=8.06E-04) as well as mitochondrial dysfunction (p=1.05E-02). Neurons rely on mitochondrial function to establish membrane excitability, neurotransmission, plasticity, and axon guidance (Kann and Kovács, 2007; Smith and Gallo, 2018). For that reason, mitochondrial alterations could affect neural maturation, neurite outgrowth, and neurotransmitter release (Levy et al., 2003; Nicholls and Budd, 2000). Studies conducted on neurons derived from schizophrenia-patient iPSCs showed reduced neuronal connectivity, a decreased number of neurites, and impaired ability to differentiate into dopaminergic and mature glutamatergic neurons (Brennand et al., 2011a; Robicsek et al., 2013). Also, by comparing the proteins found altered in the mitochondrial proteome of neurons with proteomics findings of postmortem brain tissue from schizophrenia patients (Zuccoli et al., 2017), we could identify similarities between alterations found in a model of neurodevelopment and the adult brain of schizophrenia patients.

In the total proteome of neurons, differentially regulated proteins (Sup. Table 3) were found to be related to glycolysis, TCA cycle, response to reactive oxygen species, and neuron projection development. The canonical pathways axonal guidance signaling (p=6.87E-09) and glycolysis (p=1.72E-09) were significantly altered (Figure 4B). The metabolic transition from glycolytic to oxidative (Bélanger et al., 2011) is tightly coupled to neuronal differentiation, as it was observed by decreased expression levels of glycolytic proteins. Furthermore, the constitutive expression of glycolytic enzymes during differentiation leads to neuronal cell death, indicating the decrease in aerobic glycolysis is essential for neuronal survival (Zheng et al., 2016). Our analysis revealed that glycolytic proteins were mostly upregulated in schizophrenia derived neurons when compared to controls. This indicates that the process of neuronal differentiation and, therefore, neurodevelopment could be affected in schizophrenia.

Additionally, we have found proteins related to response to ROS altered in schizophrenia, such as superoxide dismutase, peroxiredoxins, and thioredoxin. Oxidative stress is related to risk factors for neurodevelopmental disorders (Do et al., 2009) and changes in the redox system have been identified in diverse models of schizophrenia (Behrens et al., 2008; Pedrini et al., 2012). Our results obtained with NSCs revealed a trend of elevated levels of ROS in schizophrenia cells, which could interfere in neurodevelopmental aspects of the disorder, with similar results observed in other patient-derived NPCs (Paulsen et al., 2012). Hence, in the developing embryo, the occurrence of oxidative stress may affect development by damaging lipids, DNA, and proteins, potentially altering signal transduction (Wells et al., 2009) and at least partially explain the phenotypic traits. In fact, there is a link between the alterations found in the redox system and mitochondrial proteins with cytoskeleton components (Figure 4D). These alterations could interfere with the proper organization of actin filaments and microtubules, which are important for the formation and development of neuronal processes (Liu and Dwyer, 2014; Pacheco and Gallo, 2016).

## Conclusion

Considering the results discussed above, this study has led to the identification of proteins and associated pathways altered in neural cells derived from schizophrenia-patients’ iPSCs. The alterations found reveal the potential targets involved with compromised neurodevelopment in schizophrenia, since processes such as neuronal differentiation, neuritogenesis, and axonal guidance signaling are linked to our findings. In conclusion, this work reveals the importance of combining proteomic approaches with cells derived from iPSCs from schizophrenia patients in the study of the disorder to bring insights into the mechanisms related to the neurodevelopment of schizophrenia.

## Acknowledgements

We thank Gabriela Lopes Vitória for her excellent support related to stem cell generation and culture and Ludmila S. S. Bastos for cell bank expansion. We also thank Mariana Fioramonte for her support with the mass spectrometry analysis. We thank the São Paulo Research Foundation – FAPESP (grant numbers 2016/04912-2, 2018/14666-4, 2014/21035-0, 2018/22505-0, 2015/15626-8, 2017/25588-1, 2019/00098-7 and 2018/01410-1); the Brazilian National Council for Scientific and Technological Development – CNPq and the Foundation for Research Support in the State of Rio de Janeiro (FAPERJ) for funding this work.

---

Supplementary table 1: List of proteins found altered in the mitochondrial and nuclear proteomes of NSCs

Supplementary table 2: List of proteins found altered in the mitochondrial and nuclear proteomes of neurons

Supplementary table 3: List of proteins found altered in the total proteome of neurons

