## Supplementary Table 1 for "Mitochondrial, cell cycle control and neuritogenesis alterations in an iPSC-based neurodevelopmental model for schizophrenia"

| Accession | Description | Biological Process | Cellular component | p-Value | SCZ/CTRL ratio | Fraction |
|---|---|---|---|---|---|---|
| A5YKK6 | CCR4-NOT transcription complex subunit 1 | Transcription | Nucleus | 0,0009237 | 1,595361022 | Mitochondrion |
| O00264 | Membrane-associated progesterone receptor component 1 | Cell communication | Plasma membrane | 0,0294786 | 1,777994311 | Mitochondrion |
| O43615 | Mitochondrial import inner membrane translocase subunit TIM44 | Transport | Mitochondrion | 0,0045313 | 0,431193395 | Mitochondrion |
| O43776 | Asparagine--tRNA ligase_ cytoplasmic | Protein metabolism | Cytoplasm | 0,0414845 | 0,77188705 | Mitochondrion |
| O60749 | Sorting nexin-2 | Transport | Endosome | 0,0075918 | 0,769199618 | Mitochondrion |
| O75390 | Citrate synthase_ mitochondrial | Metabolism; Energy | Mitochondrion | 0,0428382 | 0,793084926 | Mitochondrion |
| O75534 | Cold shock domain-containing protein E1 | Regulation of gene e | Cytoplasm | 0,0474248 | 3,734627541 | Mitochondrion |
| O75569 | Interferon-inducible double-stranded RNA-dependent protein kinase activator A | Regulation of nucleo | Cytoplasm | 0,0468766 | 1,733078696 | Mitochondrion |
| O75643 | U5 small nuclear ribonucleoprotein 200 kDa helicase | Regulation of nucleo | Mitochondrion; Nucleus | 0,0489081 | 1,482607097 | Mitochondrion |
| O95202 | Mitochondrial proton/calcium exchanger protein | Cell communication | Mitochondrion | 0,0119772 | 0,514952979 | Mitochondrion |
| O95573 | Long-chain-fatty-acid--CoA ligase 3 | Fatty acid metabolisı | Peroxisome; Mitochondri | 0,0035945 | 0,664969567 | Mitochondrion |
| P05062 | Fructose-bisphosphate aldolase B | Metabolism; Energy | Cytoplasm | 0,001422 | 0,763814165 | Mitochondrion |
| P05091 | Aldehyde dehydrogenase_ mitochondrial | Metabolism; Energy | Mitochondrion | 0,047609 | 1,5275866 | Mitochondrion |
| P05186 | Alkaline phosphatase_ tissue-nonspecific isozyme | Metabolism; Energy | Plasma membrane | 0,0343502 | 0,464951135 | Mitochondrion |
| P06576 | ATP synthase subunit beta_ mitochondrial | Metabolism; Energy | Mitochondrion | 0,0001346 | 0,7809306 | Mitochondrion |
| P08133 | Annexin A6 | Cell communication; | Endoplasmic reticulum; N | 0,002184 | 0,324697452 | Mitochondrion |
| P08559 | Pyruvate dehydrogenase E1 component subunit alpha_ somatic form_ mitochondrial | Metabolism; Energy | Mitochondrion | 0,0001175 | 0,790939034 | Mitochondrion |
| P09110 | 3-ketoacyl-CoA thiolase_ peroxisomal | Metabolism; Energy | Peroxisome; Mitochondri | 0,0185633 | 0,61827695 | Mitochondrion |
| P12268 | Inine-5'-monophosphate dehydrogenase 2 | Metabolism; Energy | Cytoplasm | 0,0150328 | 0,547608564 | Mitochondrion |
| P12814 | Alpha-actinin-1 | Cell growth | Cytoplasm; Mitochondrio | 0,0146274 | 0,766394171 | Mitochondrion |
| P13591 | Neural cell adhesion molecule 1 | Cell communication | Plasma membrane | 0,0068742 | 1,575555504 | Mitochondrion |
| P13637 | Sodium/potassium-transporting ATPase subunit alpha-3 | Transport | Plasma membrane | 0,014996 | 1,758689192 | Mitochondrion |
| P13645 | Keratin_ type I cytkeletal 10 | Cell growth | Cytoplasm | 0,0403162 | 3,361262859 | Mitochondrion |
| P13674 | Prolyl 4-hydroxylase subunit alpha-1 | Metabolism; Energy | Endoplasmic reticulum | 0,0225059 | 0,67316565 | Mitochondrion |
| P14868 | Aspartate--tRNA ligase_ cytoplasmic | Metabolism; Energy | Cytoplasm | 0,0002214 | 0,713013807 | Mitochondrion |
| P16401 | Histone H1.5 | Regulation of nucleo | Mitochondrion; Nucleus | 0,0261905 | 1,855863056 | Mitochondrion |
| P18206 | Vinculin | Cell growth | Cytoplasm; Nucleus | 0,0127546 | 0,498951774 | Mitochondrion |
| P22695 | Cytochrome b-c1 complex subunit 2_ mitochondrial | Metabolism; Energy | Mitochondrion | 0,0056558 | 0,770080063 | Mitochondrion |
| P26232 | Catenin alpha-2 | Cell growth | Cytoplasm | 0,0491726 | 1,442861808 | Mitochondrion |
| P26639 | Threonine--tRNA ligase_ cytoplasmic | Metabolism; Energy | Cytoplasm; Mitochondrio | 0,0221601 | 0,509631713 | Mitochondrion |
| P31930 | Cytochrome b-c1 complex subunit 1_ mitochondrial | Metabolism; Energy | Mitochondrion | 0,0166592 | 0,443311936 | Mitochondrion |
| P32322 | Pyrroline-5-carboxylate reductase 1_ mitochondrial | Metabolism; Energy | Cytoplasm; Mitochondrio | 0,0036904 | 0,564038754 | Mitochondrion |
| P35637 | RNA-binding protein FUS | RNA localization | Mitochondrion; Nucleus | 0,0360421 | 1,304864405 | Mitochondrion |
| P36542 | ATP synthase subunit gamma_ mitochondrial | Metabolism; Energy | Mitochondrion | 0,0078833 | 0,825763421 | Mitochondrion |
| P37837 | Transaldolase | Metabolism; Energy | Nucleus | 0,0222407 | 4,88239712 | Mitochondrion |
| P38646 | Stress-70 protein_ mitochondrial | Protein metabolism | Mitochondrion | 0,0151956 | 0,628961503 | Mitochondrion |
| P40926 | Malate dehydrogenase_ mitochondrial | Metabolism; Energy | Mitochondrion | 0,0401738 | 0,562047494 | Mitochondrion |
| P41091 | Eukaryotic translation initiation factor 2 subunit 3 | Protein metabolism | Cytoplasm | 0,0238845 | 1,836136991 | Mitochondrion |
| P42696 | RNA-binding protein 34 | Regulation of nucleo | Nucleus | 0,0360053 | 0,841860042 | Mitochondrion |
| P42765 | 3-ketoacyl-CoA thiolase_ mitochondrial | Metabolism; Energy | Mitochondrion | 0,0027423 | 0,496185903 | Mitochondrion |
| P50454 | Serpin H1 | Protein metabolism | Endoplasmic reticulum | 0,0264448 | 0,545857348 | Mitochondrion |
| P54136 | Arginine--tRNA ligase_ cytoplasmic | Protein metabolism | Cytoplasm | 0,0002228 | 0,514482809 | Mitochondrion |
| P54886 | Delta-1-pyrroline-5-carboxylate synthase | Metabolism; Energy | Mitochondrion | 0,0144135 | 0,657968022 | Mitochondrion |
| P55010 | Eukaryotic translation initiation factor 5 | Protein metabolism | Cytoplasm | 0,0383736 | 0,733990998 | Mitochondrion |
| P55011 | Solute carrier family 12 member 2 | Transport | Plasma membrane | 0,0111084 | 1,183295731 | Mitochondrion |
| P61221 | ATP-binding cassette sub-family E member 1 | Transport | Cytoplasm; Mitochondrio | 0,0134555 | 0,791770593 | Mitochondrion |
| Q00325 | Phosphate carrier protein_ mitochondrial | Transport | Mitochondrion | 0,0205585 | 0,74524321 | Mitochondrion |
| Q02246 | Contactin-2 | Cell communication | Plasma membrane | 0,0489403 | 2,218740561 | Mitochondrion |
| Q04637 | Eukaryotic translation initiation factor 4 gamma 1 | Protein metabolism | Cytoplasm | 0,0041499 | 1,80805722 | Mitochondrion |
| Q07666 | KH domain-containing_ RNA-binding_ signal transduction-associated protein 1 | Regulation of nucleo | Nucleus | 0,0191996 | 0,751173806 | Mitochondrion |
| Q08211 | ATP-dependent RNA helicase A | Regulation of nucleo | Nucleus | 0,022207 | 1,765286881 | Mitochondrion |
| Q08945 | FACT complex subunit SSRP1 | Regulation of nucleo | Nucleus | 0,0009606 | 0,428124422 | Mitochondrion |
| Q13867 | Bleomycin hydrolase | Metabolism; Energy | Cytoplasm | 0,0178838 | 1,879747103 | Mitochondrion |
| Q15758 | Neutral amino acid transporter B(0) | Transport | Plasma membrane | 0,0486491 | 0,674360422 | Mitochondrion |
| Q16891 | MICOS complex subunit MIC60 | Cell growth | Mitochondrion | 0,0486491 | 0,583447916 | Mitochondrion |
| Q2VWP7 | Protogenin | | Plasma membrane | 0,015688 | 0,363221456 | Mitochondrion |
| Q53H12 | Acylglycerol kinase_ mitochondrial | Cell communication | Mitochondrion | 0,0413518 | 0,380615229 | Mitochondrion |
| Q5IJ48 | Protein crumbs homolog 2 | Cell growth | Plasma membrane | 4,555E-05 | 2,143476668 | Mitochondrion |
| Q5T9A4 | ATPase family AAA domain-containing protein 3A | Metabolism; Energy | Mitochondrion | 0,001546 | 0,502840027 | Mitochondrion |
| Q6UB35 | Monofunctional C1-tetrahydrofolate synthase_ mitochondrial | Metabolism | Mitochondrion | 0,0011758 | 0,627346579 | Mitochondrion |
| Q8WVM8 | Sec1 family domain-containing protein 1 | Transport | Plasma membrane | 0,0078008 | 0,803623286 | Mitochondrion |
| Q92692 | Nectin-2 | Cell communication | Plasma membrane | 0,0047758 | 0,523042969 | Mitochondrion |
| Q92896 | Golgi apparatus protein 1 | | Golgi apparatus; Mitocho | 0,0274554 | 0,764505822 | Mitochondrion |
| Q92900 | Regulator of nonsense transcripts 1 | Regulation of transla | Cytoplasm | 0,0114718 | 1,55156945 | Mitochondrion |
| Q96A65 | Exocyst complex component 4 | Transport | Plasma membrane | 0,0070869 | 1,810996959 | Mitochondrion |
| Q96AE4 | Far upstream element-binding protein 1 | Regulation of nucleo | Nucleus | 0,0067549 | 0,583002797 | Mitochondrion |
| Q96JA1 | Leucine-rich repeats and immunoglobulin-like domains protein 1 | Cell communication | Plasma membrane | 0,0390648 | 0,528261854 | Mitochondrion |
| Q96L92 | Sorting nexin-27 | Vesicle-mediated tra | Cytoplasm | 0,0494936 | 0,708765926 | Mitochondrion |
| Q96PQ0 | VPS10 domain-containing receptor SorCS2 | Cell communication | Plasma membrane | 0,0076851 | 4,348269313 | Mitochondrion |
| Q9BSJ8 | Extended synaptotagmin-1 | Cell communication | Mitochondrion; Plasma m | 0,0283158 | 0,676389776 | Mitochondrion |
| Q9H2U1 | ATP-dependent RNA helicase DHX36 | Regulation of nucleo | Plasma membrane | 0,0445643 | 0,594746642 | Mitochondrion |
| Q9H9A5 | CCR4-NOT transcription complex subunit 10 | | | 0,0152181 | 1,288597547 | Mitochondrion |
| Q9H9B4 | Sideroflexin-1 | Transport | Mitochondrion | 0,0266244 | 0,418704228 | Mitochondrion |
| Q9NYU2 | UDP-glucose:glycoprotein glucyltransferase 1 | Metabolism; Energy | Endoplasmic reticulum; N | 0,0225234 | 1,927624966 | Mitochondrion |
| Q9UH03 | Neuronal-specific septin-3 | Cell communication | Plasma membrane | 0,0368229 | 1,726913948 | Mitochondrion |
| Q9UHB9 | Signal recognition particle subunit SRP68 | Protein metabolism | Nucleus | 0,0091205 | 0,758969829 | Mitochondrion |
| Q9UKV8 | Protein argonaute-2 | Protein metabolism | Cytoplasm | 0,0219266 | 2,285783341 | Mitochondrion |
| Q9UQE7 | Structural maintenance of chromosomes protein 3 | DNA repair | Mitochondrion; Nucleus | 0,0117422 | 1,541625914 | Mitochondrion |
| Q9Y4W6 | AFG3-like protein 2 | Transport | Mitochondrion | 0,006184 | 1,249977102 | Mitochondrion |
| P11177 | Pyruvate dehydrogenase E1 component subunit beta_ mitochondrial | Metabolism; Energy | Mitochondrion | 0,0021535 | 0,737384348 | Nucleus |
| P47897 | 3-ketoacyl-CoA thiolase_ mitochondrial | Metabolism; Energy | Cytoplasm | 0,0214663 | 0,76349697 | Nucleus |
| P55084 | Trifunctional enzyme subunit beta_ mitochondrial | Metabolism; Energy | Mitochondrion | 0,0024346 | 0,80172129 | Nucleus |
| P56134 | ATP synthase subunit f_ mitochondrial | Electron transport | Mitochondrion | 0,002477 | 0,573219618 | Nucleus |
| Q9UDR5 | Alpha-aminoadipic semialdehyde synthase_ mitochondrial | Metabolism; Energy | Mitochondrion | 0,002785 | 1,443959595 | Nucleus |
| O00217 | NADH dehydrogenase [ubiquinone] iron-sulfur protein 8_ mitochondrial | Metabolism; Energy | Mitochondrion; Nucleus | 0,0306479 | 0,605321736 | Nucleus |
| O00231 | 26S proteasome non-ATPase regulatory subunit 11 | Protein metabolism | Nucleus | 0,0043536 | 1,281018966 | Nucleus |
| O00299 | Chloride intracellular channel protein 1 | Transport | Nucleus | 0,021777 | 0,756513069 | Nucleus |
| O00410 | Importin-5 | Transport | Cytoplasm; Nucleus | 0,0424261 | 1,440367442 | Nucleus |
| O00487 | 26S proteasome non-ATPase regulatory subunit 14 | Protein metabolism | Cytoplasm; Nucleus | 0,0120854 | 0,844216146 | Nucleus |
| O14579 | Coatomer subunit epsilon | Transport | Cytoplasm | 0,017233 | 0,753035955 | Nucleus |
| O14980 | Exportin-1 | Cell communication | Nucleus | 0,0051921 | 0,722022511 | Nucleus |
| O15042 | U2 snRNP-associated SURP motif-containing protein | RNA binding | Nucleus | 0,0017928 | 1,76349104 | Nucleus |
| O15523 | ATP-dependent RNA helicase DDX3Y | Regulation of nucleo | Nucleus | 0,001938 | 1,444579811 | Nucleus |
| O43347 | RNA-binding protein Musashi homolog 1 | Regulation of nucleo | Cytoplasm | 0,0174603 | 1,31910785 | Nucleus |
| O43399 | Tumor protein D54 | Cell proliferation | Cytoplasm | 0,0020641 | 0,872310662 | Nucleus |

| | | | | | | |
|---|---|---|---|---|---|---|
| O60664 | Perilipin-3 | Transport | Cytoplasm | 0,0056275 | 0,64528828 | Nucleus |
| O60716 | Catenin delta-1 | Cell communication | Nucleus | 0,0352407 | 0,796004824 | Nucleus |
| O75376 | Nuclear receptor corepressor 1 | Regulation of nucleo | Nucleus | 0,0074645 | 0,661967125 | Nucleus |
| O75400 | pre-mRNA processing factor 40 homolog A | Regulation of nucleo | Nucleus | 0,0204413 | 1,446789233 | Nucleus |
| O75521 | Enoyl-CoA delta isomerase 2_ mitochondrial | Fatty acid metabolisı | Peroxisome | 0,0009043 | 0,59778388 | Nucleus |
| O75643 | U5 small nuclear ribonucleoprotein 200 kDa helicase | Regulation of nucleo | Mitochondrion; Nucleus | 0,0055332 | 1,842977912 | Nucleus |
| O75874 | Isocitrate dehydrogenase [NADP] cytoplasmic | Metabolism; Energy | Cytoplasm | 0,0046762 | 1,472463678 | Nucleus |
| O76094 | Signal recognition particle subunit SRP72 | Protein metabolism | Nucleus | 0,0007327 | 0,785876404 | Nucleus |
| O95292 | Vesicle-associated membrane protein-associated protein B/C | Transport | Plasma membrane | 0,0107871 | 0,768013028 | Nucleus |
| O95373 | Importin-7 | Transport | Nucleus | 0,0002003 | 1,929225039 | Nucleus |
| O95831 | Apoptis-inducing factor 1_ mitochondrial | Cell communication | Mitochondrion; Nucleus | 3,864E-06 | 0,729113552 | Nucleus |
| O95865 | N(G)_N(G)-dimethylarginine dimethylaminohydrolase 2 | Metabolism; Energy | Cytoplasm | 0,0064616 | 1,481259417 | Nucleus |
| O96008 | Mitochondrial import receptor subunit TOM40 homolog | Transport | Mitochondrion | 0,0067282 | 0,784513783 | Nucleus |
| O96019 | Actin-like protein 6A | Regulation of nucleo | Nucleus | 0,0014446 | 1,222279174 | Nucleus |
| P02768 | Serum albumin | | | 0,0137512 | 1,967135865 | Nucleus |
| P05186 | Alkaline phosphatase_ tissue-nonspecific isozyme | Metabolism; Energy | Plasma membrane | 0,0489548 | 0,654321155 | Nucleus |
| P05388 | 60S acidic ribomal protein P0 | Protein metabolism | Nucleus; Ribosome | 4,641E-05 | 0,873161677 | Nucleus |
| P06493 | Cyclin-dependent kinase 1 | Cell communication | Nucleus | 0,0382183 | 1,24979603 | Nucleus |
| P06748 | Nucleophosmin | Protein metabolism | Nucleus | 0,012087 | 0,718519823 | Nucleus |
| P07195 | L-lactate dehydrogenase B chain | Metabolism; Energy | Cytoplasm | 0,0100615 | 1,240578646 | Nucleus |
| P07196 | Neurofilament light polypeptide | Cell growth | Cytoplasm | 0,0253412 | 0,761379621 | Nucleus |
| P07305 | Histone H1.0 | Regulation of nucleo | Nucleus | 0,0155675 | 0,414628093 | Nucleus |
| P07355 | Annexin A2 | Cell communication; | Nucleus | 0,0199679 | 0,386828239 | Nucleus |
| P08133 | Annexin A6 | Cell communication; | Endoplasmic reticulum | 0,0063335 | 0,533839253 | Nucleus |
| P08195 | 4F2 cell-surface antigen heavy chain | Transport | Plasma membrane | 0,0071986 | 0,741756518 | Nucleus |
| P09211 | Glutathione S-transferase P | Metabolism; Energy | Cytoplasm | 0,0066129 | 1,441695885 | Nucleus |
| P09874 | Poly [ADP-ribe] polymerase 1 | Protein metabolism | Nucleus | 0,0233723 | 1,839690387 | Nucleus |
| P09936 | Ubiquitin carboxyl-terminal hydrolase isozyme L1 | Protein metabolism | Cytoplasm | 0,0199742 | 1,349334094 | Nucleus |
| P0DP23 | Calmodulin-1 | Cell communication | Cytoplasm; Nucleus | 0,0173871 | 2,059206212 | Nucleus |
| P10809 | 60 kDa heat shock protein_ mitochondrial | Apoptosis | Cytoplasm; Mitochondrio | 0,0121135 | 0,737504613 | Nucleus |
| P11137 | Microtubule-associated protein 2 | Cell growth | Cytoplasm | 0,0405392 | 1,391334025 | Nucleus |
| P11216 | Glycogen phosphorylase_ brain form | Metabolism; Energy | Cytoplasm; Nucleus | 0,0127037 | 1,54782519 | Nucleus |
| P11279 | Lysosome-associated membrane glycoprotein 1 | | Lysossome | 0,0023436 | 1,91244341 | Nucleus |
| P11766 | Alcohol dehydrogenase class-3 | Metabolism; Energy | Cytoplasm; Nucleus | 0,0272111 | 1,262381341 | Nucleus |
| P12277 | Creatine kinase B-type | Metabolism; Energy | Cytoplasm | 0,0235189 | 1,340169861 | Nucleus |
| P12956 | X-ray repair crs-complementing protein 6 | Regulation of nucleo | Nucleus | 0,0359676 | 1,286150284 | Nucleus |
| P13010 | X-ray repair crs-complementing protein 5 | Regulation of nucleo | Nucleus | 0,0351366 | 1,416605281 | Nucleus |
| P13645 | Keratin_ type I cytoskeletal 10 | Cell growth | Cytoplasm | 0,0006093 | 1,918803251 | Nucleus |
| P15121 | Aldose reductase | Metabolism; Energy | Cytoplasm | 0,0004901 | 1,557729445 | Nucleus |
| P16401 | Histone H1.5 | Regulation of nucleo | Nucleus | 0,000231 | 3,302855205 | Nucleus |
| P18206 | Vinculin | Cell growth | Cytoplasm; Nucleus | 0,0483305 | 0,835594344 | Nucleus |
| P18754 | Regulator of chromosome condensation | Chromosome organi: | Nucleus | 0,0313523 | 1,852513166 | Nucleus |
| P19022 | Cadherin-2 | Cell communication | Plasma membrane | 0,0020416 | 1,588244438 | Nucleus |
| P21796 | Voltage-dependent anion-selective channel protein 1 | Transport | Mitochondrion; Nucleus | 0,0012501 | 0,748755784 | Nucleus |
| P22570 | NADPH:adrenodoxin oxidoreductase_ mitochondrial | Metabolism; Energy | Mitochondrion | 0,0003428 | 0,454797616 | Nucleus |
| P23193 | Transcription elongation factor A protein 1 | Transcription regulat | Nucleus | 0,0058946 | 1,294542341 | Nucleus |
| P24752 | Acetyl-CoA acetyltransferase_ mitochondrial | Metabolism; Energy | Cytoplasm; Mitochondrio | 0,0072783 | 0,578772433 | Nucleus |
| P24941 | Cyclin-dependent kinase 2 | Regulation of nucleo | Nucleus | 0,0495074 | 1,544028849 | Nucleus |
| P25205 | DNA replication licensing factor MCM3 | Regulation of nucleo | Nucleus | 0,0078507 | 1,739156286 | Nucleus |
| P25705 | ATP synthase subunit alpha_ mitochondrial | Metabolism; Energy | Mitochondrion | 9,23E-05 | 0,827332138 | Nucleus |
| P26232 | Catenin alpha-2 | Cell growth | Cytoplasm | 0,0236046 | 0,751298556 | Nucleus |
| P27348 | 14-3-3 protein theta | Cell communication | Cytoplasm; Nucleus | 0,0079458 | 1,296587065 | Nucleus |
| P27694 | Replication protein A 70 kDa DNA-binding subunit | Regulation of nucleo | Nucleus | 0,0487831 | 1,286814621 | Nucleus |
| P27816 | Microtubule-associated protein 4 | Cell growth | Cytoplasm | 0,0076435 | 1,700514421 | Nucleus |
| P29762 | Cellular retinoic acid-binding protein 1 | Cell growth | Cytoplasm | 0,0127374 | 0,605029271 | Nucleus |
| P29966 | Myristoylated alanine-rich C-kinase substrate | Cell growth | Plasma membrane | 0,006199 | 1,474818866 | Nucleus |
| P30086 | Phosphatidylethanolamine-binding protein 1 | Cell communication | Cytoplasm | 0,0099211 | 1,694175893 | Nucleus |
| P30154 | Serine/threonine-protein phphatase 2A 65 kDa regulatory subunit A beta isoform | Cell communication | Cytoplasm; Nucleus | 0,0348075 | 0,779649087 | Nucleus |
| P30566 | Adenyluccinate lyase | Metabolism; Energy | Cytoplasm | 0,000569 | 0,650299071 | Nucleus |
| P30837 | Aldehyde dehydrogenase X_ mitochondrial | Metabolism; Energy | Mitochondrion | 0,0355562 | 0,697917353 | Nucleus |
| P31150 | Rab GDP dissociation inhibitor alpha | Cell communication | Cytoplasm | 0,0085507 | 1,461240665 | Nucleus |
| P31153 | S-adenylmethionine synthase isoform type-2 | Metabolism; Energy | Cytoplasm | 0,0143085 | 1,300154834 | Nucleus |
| P31689 | DnaJ homolog subfamily A member 1 | Protein metabolism | Nucleus | 0,030291 | 0,808453948 | Nucleus |
| P32119 | Peroxiredoxin-2 | Metabolism; Energy | Cytoplasm | 0,003068 | 1,579909952 | Nucleus |
| P33991 | DNA replication licensing factor MCM4 | Regulation of nucleo | Nucleus | 0,0006213 | 1,891175997 | Nucleus |
| P34932 | Heat shock 70 kDa protein 4 | Protein metabolism | Cytoplasm; Nucleus | 0,0037123 | 1,464130294 | Nucleus |
| P35221 | Catenin alpha-1 | Cell growth | Cytoplasm | 0,0280463 | 1,250977204 | Nucleus |
| P35232 | Prohibitin | Cell communication | Nucleus; Mitochondrion | 5,583E-05 | 0,699925527 | Nucleus |
| P35241 | Radixin | Cell growth | Plasma membrane | 0,0404667 | 1,516744115 | Nucleus |
| P35580 | Myosin-10 | Cell growth | Cytoplasm | 0,0002724 | 2,899764052 | Nucleus |
| P35611 | Alpha-adducin | Cell growth | Nucleus | 0,0254077 | 1,885491379 | Nucleus |
| P35613 | Basigin | Cell communication | Plasma membrane | 0,013047 | 0,772952753 | Nucleus |
| P36507 | Dual specificity mitogen-activated protein kinase kinase 2 | Cell communication | Cytoplasm; Nucleus | 0,0356278 | 0,706223679 | Nucleus |
| P37837 | Transaldolase | Metabolism; Energy | Nucleus | 0,0152573 | 1,385335869 | Nucleus |
| P38646 | Stress-70 protein_ mitochondrial | Protein metabolism | Mitochondrion; Nucleus | 0,0063581 | 0,731543193 | Nucleus |
| P40925 | Malate dehydrogenase_ cytoplasmic | Metabolism; Energy | Cytoplasm; Mitochondrio | 0,0211028 | 1,329279095 | Nucleus |
| P40926 | Malate dehydrogenase_ mitochondrial | Metabolism; Energy | Mitochondrion; Nucleus | 0,0001451 | 0,697438257 | Nucleus |
| P40937 | Replication factor C subunit 5 | DNA replication | Nucleus | 0,0037681 | 1,338074733 | Nucleus |
| P40939 | Trifunctional enzyme subunit alpha_ mitochondrial | Metabolism; Energy | Mitochondrion; Nucleus | 9,028E-05 | 0,699501609 | Nucleus |
| P43243 | Matrin-3 | Regulation of nucleo | Nucleus | 0,0065288 | 1,543260217 | Nucleus |
| P43246 | DNA mismatch repair protein Msh2 | Regulation of nucleo | Nucleus | 0,0249427 | 1,278087689 | Nucleus |
| P43307 | Translocon-associated protein subunit alpha | Transport | Endoplasmic reticulum | 0,0013172 | 0,62395124 | Nucleus |
| P45880 | Voltage-dependent anion-selective channel protein 2 | Transport | Mitochondrion; Nucleus | 0,0043666 | 0,757043367 | Nucleus |
| P46821 | Microtubule-associated protein 1B | Cell growth | Cytoplasm; Nucleus | 8,784E-05 | 1,424403531 | Nucleus |
| P47985 | Putative cytochrome b-c1 complex subunit Rieske-like protein 1 | Metabolism; Energy | Mitochondrion | 0,0131561 | 0,804110554 | Nucleus |
| P48681 | Nestin | Cell growth | Cytoplasm; Nucleus | 0,0068516 | 1,372493825 | Nucleus |
| P49321 | Nuclear autoantigenic sperm protein | Cell communication | Nucleus | 0,0164462 | 1,741410424 | Nucleus |
| P49411 | Elongation factor Tu_ mitochondrial | Protein metabolism | Mitochondrion; Nucleus | 0,0040793 | 0,748386235 | Nucleus |
| P49721 | Proteasome subunit beta type-2 | Protein metabolism | Cytoplasm | 0,0137118 | 1,238550987 | Nucleus |
| P49736 | DNA replication licensing factor MCM2 | Regulation of nucleo | Nucleus | 6,134E-05 | 2,899562015 | Nucleus |
| P49821 | NADH dehydrogenase [ubiquinone] flavoprotein 1_ mitochondrial | Metabolism; Energy | Mitochondrion | 0,0002183 | 0,750661135 | Nucleus |
| P50402 | Emerin | Cell growth | Nucleus | 0,0010072 | 0,67209495 | Nucleus |
| P50454 | Serpin H1 | Protein metabolism | Endoplasmic reticulum | 0,0021153 | 0,480225844 | Nucleus |
| P51398 | 28S ribomal protein S29_ mitochondrial | Apoptosis | Mitochondrion; Nucleus | 0,0234592 | 0,704110044 | Nucleus |
| P52701 | DNA mismatch repair protein Msh6 | DNA repair | Nucleus | 0,0089123 | 2,357547614 | Nucleus |
| P52735 | Guanine nucleotide exchange factor VAV2 | Cell communication | Cytoplasm; Nucleus | 0,0389437 | 0,740101524 | Nucleus |

| | | | | | | |
|---|---|---|---|---|---|---|
| P53396 | ATP-citrate synthase | Metabolism; Energy | Cytoplasm; Nucleus | 0,0007074 | 1,356341786 | Nucleus |
| P53597 | Succinate--CoA ligase [ADP/GDP-forming] subunit alpha_ mitochondrial | Metabolism; Energy | Mitochondrion | 0,0044326 | 0,789923219 | Nucleus |
| P53990 | IST1 homolog | Cell division | Nucleus | 0,0167539 | 0,827505198 | Nucleus |
| P54709 | Sodium/potassium-transporting ATPase subunit beta-3 | Transport | Plasma membrane | 0,0041801 | 0,848523462 | Nucleus |
| P54886 | Delta-1-pyrroline-5-carboxylate synthase | Metabolism; Energy | Mitochondrion | 0,0238918 | 0,826681518 | Nucleus |
| P55786 | Puromycin-sensitive aminopeptidase | Protein metabolism | Cytoplasm; Nucleus | 0,0096514 | 1,435377913 | Nucleus |
| P60174 | Triosephosphate isomerase | Metabolism; Energy | Cytoplasm | 0,0283189 | 1,253797993 | Nucleus |
| P60900 | Proteasome subunit alpha type-6 | Protein metabolism | Cytoplasm; Nucleus | 0,0461687 | 1,097990517 | Nucleus |
| P61254 | 60S ribomal protein L26-like 1 | Protein metabolism | Ribosome | 0,0136037 | 1,276488666 | Nucleus |
| P61604 | 10 kDa heat shock protein_ mitochondrial | Protein metabolism | Mitochondrion | 0,0145687 | 0,456229448 | Nucleus |
| P62136 | Serine/threonine-protein phosphatase PP1-alpha catalytic subunit | Cell proliferation | Cytoeskeleton | 0,0108403 | 0,888525723 | Nucleus |
| P62280 | 40S ribosomal protein S11 | Protein metabolism | Ribosome; Nucleus | 0,0009815 | 1,468008271 | Nucleus |
| P62333 | 26S proteasome regulatory subunit 10B | Protein metabolism | Cytoplasm | 0,0150883 | 0,881464707 | Nucleus |
| P62495 | Eukaryotic peptide chain release factor subunit 1 | Protein metabolism | Ribosome; Nucleus | 2,602E-05 | 0,842095085 | Nucleus |
| P62906 | 60S ribosomal protein L10a | Protein metabolism | Ribosome; Nucleus | 0,0110262 | 0,82339368 | Nucleus |
| P62913 | 60S ribosomal protein L11 | Protein metabolism | Ribosome; Nucleus | 0,01699 | 1,40601326 | Nucleus |
| P63104 | 14-3-3 protein zeta/delta | Regulation of cell cyc | Cytoplasm | 0,026998 | 1,237409863 | Nucleus |
| P78344 | Eukaryotic translation initiation factor 4 gamma 2 | Protein metabolism | Cytoplasm | 0,0105219 | 0,875849295 | Nucleus |
| P78527 | DNA-dependent protein kinase catalytic subunit | Cell communication | Nucleus | 0,0097186 | 2,155032306 | Nucleus |
| P98172 | Ephrin-B1 | Cell communication | Plasma membrane | 0,0285825 | 0,631650926 | Nucleus |
| Q00341 | Vigilin | Transport | Cytoplasm | 0,0266754 | 1,425695984 | Nucleus |
| Q01581 | Hydroxymethylglutaryl-CoA synthase_ cytoplasmic | Metabolism; Energy | Cytoplasm | 0,0070479 | 1,540424666 | Nucleus |
| Q02880 | DNA topoisomerase 2-beta | Regulation of nucleo | Nucleus | 0,0208909 | 2,206810337 | Nucleus |
| Q04760 | Lactoylglutathione lyase | Metabolism; Energy | Cytoplasm | 0,0015114 | 1,373978859 | Nucleus |
| Q04917 | 14-3-3 protein eta | Cell communication | Cytoplasm; Nucleus | 0,0085051 | 1,239102838 | Nucleus |
| Q12874 | Splicing factor 3A subunit 3 | Protein metabolism | Nucleus | 0,0158199 | 1,145014629 | Nucleus |
| Q12931 | Heat shock protein 75 kDa_ mitochondrial | Protein metabolism | Mitochondrion | 0,0005252 | 0,697217074 | Nucleus |
| Q13084 | 39S ribomal protein L28_ mitochondrial | Protein metabolism | Mitochondrion | 0,0060977 | 0,840295686 | Nucleus |
| Q13148 | TAR DNA-binding protein 43 | Regulation of nucleo | Nucleus | 3,507E-05 | 1,186731352 | Nucleus |
| Q13247 | Serine/arginine-rich splicing factor 6 | Regulation of nucleo | Nucleus | 0,0015214 | 1,327529017 | Nucleus |
| Q14194 | Dihydropyrimidinase-related protein 1 | Cell growth | Cytoplasm | 0,0009313 | 1,206753794 | Nucleus |
| Q14195 | Dihydropyrimidinase-related protein 3 | Metabolism; Energy | Cytoplasm; Nucleus | 0,0006223 | 1,602860636 | Nucleus |
| Q14204 | Cytoplasmic dynein 1 heavy chain 1 | Metabolism; Energy | Cytoplasm; Nucleus | 0,0059347 | 1,887803685 | Nucleus |
| Q14974 | Importin subunit beta-1 | Transport | Cytoplasm; Nucleus | 0,0255627 | 1,190805069 | Nucleus |
| Q14980 | Nuclear mitotic apparatus protein 1 | Cell growth | Nucleus | 0,0120603 | 1,649404355 | Nucleus |
| Q15046 | Lysine--tRNA ligase | tRNA aminoacylation | Cytoplasm | 0,03484 | 0,464919535 | Nucleus |
| Q15050 | Ribosome biogenesis regulatory protein homolog | Cell growth | Nucleus | 0,0184416 | 0,664757067 | Nucleus |
| Q15758 | Neutral amino acid transporter B(0) | Transport | Plasma membrane | 0,0327203 | 0,796822203 | Nucleus |
| Q15785 | Mitochondrial import receptor subunit TOM34 | Transport | Cytoplasm; Mitochondrio | 0,0093709 | 0,630439497 | Nucleus |
| Q16352 | Alpha-internexin | Cell growth | Cytoplasm | 0,030307 | 1,329020331 | Nucleus |
| Q16555 | Dihydropyrimidinase-related protein 2 | Cell communication | Cytoplasm; Nucleus | 0,0014631 | 1,474487689 | Nucleus |
| Q16658 | Fascin | Cell growth | Cytoplasm | 0,03986 | 1,309283779 | Nucleus |
| Q16698 | 2_4-dienoyl-CoA reductase_ mitochondrial | Metabolism; Energy | Mitochondrion | 0,0007467 | 0,602903539 | Nucleus |
| Q16795 | NADH dehydrogenase [ubiquinone] 1 alpha subcomplex subunit 9_ mitochondrial | Metabolism; Energy | Mitochondrion | 0,0035356 | 0,794531574 | Nucleus |
| Q16836 | Hydroxyacyl-coenzyme A dehydrogenase_ mitochondrial | | Mitochondrion | 0,0001593 | 0,728036577 | Nucleus |
| Q3ZCQ8 | Mitochondrial import inner membrane translocase subunit TIM50 | Cell communication | Mitochondrion | 0,0040656 | 0,767693723 | Nucleus |
| Q53GS9 | U4/U6.U5 tri-snRNP-associated protein 2 | Protein metabolism | Nucleus | 0,0135297 | 1,214631464 | Nucleus |
| Q5JTV8 | Torsin-1A-interacting protein 1 | | Nucleus | 0,0175815 | 1,232698256 | Nucleus |
| Q5SSJ5 | Heterochromatin protein 1-binding protein 3 | Regulation of nucleo | Nucleus | 0,0192131 | 1,375047374 | Nucleus |
| Q6NUK1 | Calcium-binding mitochondrial carrier protein SCaMC-1 | Transport | Mitochondrion | 0,0132478 | 0,668198619 | Nucleus |
| Q6P2Q9 | Pre-mRNA-processing-splicing factor 8 | Regulation of nucleo | Nucleus | 0,0128968 | 4,755357163 | Nucleus |
| Q6YN16 | Hydroxysteroid dehydrogenase-like protein 2 | | | 0,0236716 | 0,680714392 | Nucleus |
| Q7Z434 | Mitochondrial antiviral-signaling protein | | | 0,0016091 | 0,633838308 | Nucleus |
| Q7Z7K6 | Centromere protein V | Regulation of nucleo | Nucleus | 0,0023226 | 1,831354051 | Nucleus |
| Q8N0Y7 | Probable phosphoglycerate mutase 4 | Metabolism | | 0,0086863 | 1,246332337 | Nucleus |
| Q8N163 | Cell cycle and apoptis regulator protein 2 | Signal transduction | Nucleus | 0,0137423 | 1,406215794 | Nucleus |
| Q8N1F7 | Nuclear pore complex protein Nup93 | Transport | Nucleus | 0,0354165 | 0,847694533 | Nucleus |
| Q8N684 | Cleavage and polyadenylation specificity factor subunit 7 | RNA binding | Nucleus | 0,0002926 | 1,133214292 | Nucleus |
| Q8NBS9 | Thioredoxin domain-containing protein 5 | Protein metabolism | Endoplasmic reticulum | 0,0333698 | 0,759042926 | Nucleus |
| Q8NHW5 | 60S acidic ribomal protein P0-like | Ribosome biogenesis | Ribosome | 0,0025482 | 0,87203174 | Nucleus |
| Q8TAQ2 | SWI/SNF complex subunit SMARCC2 | Regulation of nucleo | Nucleus | 3,484E-05 | 2,00947324 | Nucleus |
| Q8WXF1 | Paraspeckle component 1 | Regulation of nucleo | Nucleus | 0,0137757 | 1,115269529 | Nucleus |
| Q92552 | 28S ribomal protein S27_ mitochondrial | Protein metabolism | Mitochondrion | 0,0181253 | 0,648163702 | Nucleus |
| Q92598 | Heat shock protein 105 kDa | Protein metabolism | Cytoplasm; Nucleus | 0,0285091 | 1,107484066 | Nucleus |
| Q92618 | Zinc finger protein 516 | Regulation of nucleo | Nucleus | 0,0033606 | 0,68524731 | Nucleus |
| Q92922 | SWI/SNF complex subunit SMARCC1 | Regulation of nucleo | Nucleus | 0,000721 | 1,458669976 | Nucleus |
| Q969Z0 | Protein TBRG4 | Cell communication | Mitochondrion; Nucleus | 0,0028361 | 0,741844881 | Nucleus |
| Q96A33 | Coiled-coil domain-containing protein 47 | | Mitochondrion | 0,0173569 | 0,866674419 | Nucleus |
| Q96AG4 | Leucine-rich repeat-containing protein 59 | | Plasma membrane | 0,0233578 | 0,841720393 | Nucleus |
| Q96EY1 | DnaJ homolog subfamily A member 3_ mitochondrial | Apoptosis | Mitochondrion; Nucleus | 0,0154667 | 0,526020764 | Nucleus |
| Q96EY7 | Pentatricopeptide repeat domain-containing protein 3_ mitochondrial | | Mitochondrion | 0,0069085 | 0,748401004 | Nucleus |
| Q96P70 | Importin-9 | Transport | Cytoplasm | 0,0466024 | 1,66941413 | Nucleus |
| Q96T88 | E3 ubiquitin-protein ligase UHRF1 | Regulation of nucleo | Nucleus | 0,0222728 | 1,302288276 | Nucleus |
| Q99439 | Calponin-2 | Cell growth | Cytoplasm; Nucleus | 0,018401 | 0,750387588 | Nucleus |
| Q99459 | Cell division cycle 5-like protein | Cell communication | Nucleus | 0,0002151 | 1,206470185 | Nucleus |
| Q99497 | Protein/nucleic acid deglycase DJ-1 | Regulation of nucleo | Cytoplasm; Nucleus | 0,0075854 | 1,406807462 | Nucleus |
| Q99623 | Prohibitin-2 | Regulation of nucleo | Nucleus | 0,0001981 | 0,766223336 | Nucleus |
| Q99996 | A-kinase anchor protein 9 | Cell communication | Centrosome | 0,0487522 | 0,697610123 | Nucleus |
| Q9BPU6 | Dihydropyrimidinase-related protein 5 | | Cytoplasm | 0,0287043 | 1,240979888 | Nucleus |
| Q9BPW8 | Protein NipSnap homolog 1 | | Mitochondrion | 0,0087752 | 0,830263397 | Nucleus |
| Q9BUF5 | Tubulin beta-6 chain | Cell growth | | 0,0473765 | 0,572346461 | Nucleus |
| Q9BUJ2 | Heterogeneous nuclear ribonucleoprotein U-like protein 1 | Regulation of nucleo | Nucleus | 0,0192897 | 0,690952946 | Nucleus |
| Q9BXP5 | Serrate RNA effector molecule homolog | | Nucleus | 0,0003402 | 1,30904196 | Nucleus |
| Q9BY77 | Polymerase delta-interacting protein 3 | Cell growth | Nucleus | 0,0432273 | 1,189206549 | Nucleus |
| Q9BYD2 | 39S ribomal protein L9_ mitochondrial | Protein metabolism | Mitochondrion | 0,0014386 | 0,721458906 | Nucleus |
| Q9H9B4 | Sideroflexin-1 | Transport | Mitochondrion | 0,0266244 | 0,814175432 | Nucleus |
| Q9H9Z2 | Protein lin-28 homolog A | Regulation of nucleo | Nucleus | 0,01425 | 0,528350834 | Nucleus |
| Q9NP92 | 39S ribomal protein S30_ mitochondrial | Protein metabolism | Mitochondrion | 0,0121391 | 0,525371976 | Nucleus |
| Q9NPH2 | Inositol-3-phosphate synthase 1 | Metabolism; Energy pathways | | 0,0265981 | 1,425305828 | Nucleus |
| Q9NRY4 | Rho GTPase-activating protein 35 | Cell communication | Nucleus | 0,0147827 | 1,394853418 | Nucleus |
| Q9NTX5 | Ethylmalonyl-CoA decarboxylase | | | 0,0062588 | 0,662810499 | Nucleus |
| Q9NX20 | 39S ribosomal protein L16_ mitochondrial | Protein metabolism | Mitochondrion | 0,0018707 | 0,631669254 | Nucleus |
| Q9NX40 | OCIA domain-containing protein 1 | | | 0,0005792 | 0,526127108 | Nucleus |
| Q9NZL9 | Methionine adenyltransferase 2 subunit beta | Metabolism | Cytoplasm | 0,0471674 | 1,29811784 | Nucleus |
| Q9P0L0 | Vesicle-associated membrane protein-associated protein A | Transport | Endoplasmic reticulum | 0,0017612 | 0,758079853 | Nucleus |
| Q9P2K5 | Myelin expression factor 2 | Regulation of nucleo | Nucleus | 0,0195714 | 1,157117411 | Nucleus |

| | | | | | | |
|---|---|---|---|---|---|---|
| Q9UH03 | Neuronal-specific septin-3 | Cell communication | Plasma membrane | 0,0024981 | 1,796795236 | Nucleus |
| Q9UHX1 | Poly(U)-binding-splicing factor PUF60 | Regulation of nucleo | Nucleus | 0,0353126 | 1,374046301 | Nucleus |
| Q9UI15 | Transgelin-3 | | Cytoplasm | 0,0395711 | 1,659196086 | Nucleus |
| Q9UIG0 | Tyrine-protein kinase BAZ1B | Regulation of nucleo | Nucleus | 0,0311342 | 1,846011437 | Nucleus |
| Q9UJS0 | Calcium-binding mitochondrial carrier protein Aralar2 | Transport | Mitochondrion | 6,853E-05 | 0,780551637 | Nucleus |
| Q9UJZ1 | Stomatin-like protein 2_ mitochondrial | Transport | Plasma membrane | 0,0113586 | 0,733084258 | Nucleus |
| Q9UKA9 | Polypyrimidine tract-binding protein 2 | Regulation of nucleo | Nucleus | 0,0014908 | 1,570436866 | Nucleus |
| Q9UNW9 | RNA-binding protein Nova-2 | Regulation of nucleo | Nucleus | 0,0082712 | 0,573165702 | Nucleus |
| Q9UQE7 | Structural maintenance of chromosomes protein 3 | DNA repair | Mitochondrion; Nucleus | 0,0041007 | 1,773885588 | Nucleus |
| Q9Y2W1 | Thyroid hormone receptor-associated protein 3 | Regulation of nucleo | Nucleus | 0,0469822 | 1,40556417 | Nucleus |
| Q9Y3B3 | Transmembrane emp24 domain-containing protein 7 | | | 0,002251 | 0,688811445 | Nucleus |
| Q9Y5X3 | Sorting nexin-5 | Transport | Endosome | 0,013653 | 0,493559358 | Nucleus |
