## Supplementary Table 2 for "Mitochondrial, cell cycle control and neuritogenesis alterations in an iPSC-based neurodevelopmental model for schizophrenia"

| Accession | Description | Biological Process | Cellular component | p-Value | SCZ/CTRL Ratio | Fraction |
|---|---|---|---|---|---|---|
| O00264 | Membrane-associated progesterone receptor cor | Cell communication | Plasma membrane | 0,041956435 | 1,781517241 | Mitochondrion |
| O00425 | Insulin-like growth factor 2 mRNA-binding proteir | Protein Metabolism | Cytoplasm | 0,000103789 | 3,139123808 | Mitochondrion |
| O14983 | Sarcoplasmic/endoplasmic reticulum calcium ATF | Metabolism; Energy pathways | Endoplasmic reticulum; Mitochondrion | 0,004801781 | 2,70538634 | Mitochondrion |
| O43157 | Plexin-B1 | Cell communication | Plasma membrane | 0,000381837 | 3,499203922 | Mitochondrion |
| O43295 | SLIT-ROBO Rho GTPase-activating protein 3 | Cell communication | Cytoplasm | 0,040472563 | 0,520471903 | Mitochondrion |
| O43854 | EGF-like repeat and discoidin I-like domain-contai | Cell growth | Extracellular | 0,015393309 | 0,448336632 | Mitochondrion |
| O60831 | PRA1 family protein 2 | Transport | Endosome | 7,51068E-05 | 1,716732982 | Mitochondrion |
| O75110 | Probable phospholipid-transporting ATPase IIA | Regulation of enzyme activity | Endosome | 0,000426487 | 1,993766936 | Mitochondrion |
| O75131 | Copine-3 | Transport | Cytoplasm | 0,001100983 | 5,043723328 | Mitochondrion |
| O75489 | NADH dehydrogenase [ubiquinone] iron-sulfur pr | Metabolism; Energy pathways | Mitochondrion | 3,1676E-06 | 0,574936371 | Mitochondrion |
| O75569 | Interferon-inducible double-stranded RNA-depen | Regulation of nucleotide metabolism | Cytoplasm | 0,046572176 | 1,569289619 | Mitochondrion |
| O94856 | Neurofascin | Cell communication | Plasma membrane | 0,000249579 | 1,820158646 | Mitochondrion |
| O94874 | E3 UFM1-protein ligase 1 | Protein Metabolism | Mitochondrion | 0,00028529 | 2,099486341 | Mitochondrion |
| O94979 | Protein transport protein Sec31A | Transport | Endoplasmic reticulum | 0,039789549 | 2,504856189 | Mitochondrion |
| O95336 | 6-phosphogluconolactonase | Metabolism; Energy pathways | Cytoplasm | 0,045754671 | 0,398981332 | Mitochondrion |
| O95573 | Long-chain-fatty-acid--CoA ligase 3 | Metabolism; Energy pathways | Mitochondrion; Peroxisome | 0,036001164 | 2,745340428 | Mitochondrion |
| O95865 | N(G)_N(G)-dimethylarginine dimethylaminohydro | Metabolism; Energy pathways | Cytoplasm | 0,009452516 | 2,323663442 | Mitochondrion |
| P00387 | NADH-cytochrome b5 reductase 3 | Metabolism; Energy pathways | Cytoplasm; Mitochondrion | 3,06501E-05 | 1,911969534 | Mitochondrion |
| P00558 | Phosphoglycerate kinase 1 | Metabolism; Energy pathways | Cytoplasm; Mitochondrion | 0,001831699 | 0,648253339 | Mitochondrion |
| P02768 | Serum albumin | Transport | Extracellular | 0,008786728 | 0,264157585 | Mitochondrion |
| P02786 | Transferrin receptor protein 1 | Transport | Plasma membrane; Mitochondrion | 0,00815584 | 1,684719204 | Mitochondrion |
| P04264 | Keratin_ type II cytoskeletal 1 | Cell growth | Plasma membrane | 0,039765274 | 0,272095146 | Mitochondrion |
| P04844 | Dolichyl-diphosphooligosaccharide--protein glyco | Protein Metabolism | Endoplasmic reticulum; Mitochondrion | 0,003496916 | 2,374823891 | Mitochondrion |
| P05141 | ADP/ATP translocase 2 | Metabolism; Energy pathways | Mitochondrion | 0,031299548 | 1,263580596 | Mitochondrion |
| P05455 | Lupus La protein | Regulation of nucleotide metabolism | Nucleus | 0,00147714 | 0,008726842 | Mitochondrion |
| P06733 | Alpha-enolase | Metabolism; Energy pathways | Cytoplasm | 0,001423292 | 0,669050675 | Mitochondrion |
| P07196 | Neurofilament light polypeptide | Cell growth | Cytoplasm | 0,029362404 | 0,622892521 | Mitochondrion |
| P07237 | Protein disulfide-isomerase | Protein Metabolism | Endoplasmic reticulum; Mitochondrion | 0,000224264 | 1,829078767 | Mitochondrion |
| P07437 | Tubulin beta chain | Cell growth | Cytoplasm | 0,004661952 | 0,340422923 | Mitochondrion |
| P07947 | Tyrosine-protein kinase Yes | Cell communication | Cytoplasm | 0,000671156 | 2,516998979 | Mitochondrion |
| P08133 | Annexin A6 | Cell communication | Endoplasmic reticulum; Mitochondrion | 0,023850071 | 0,623862819 | Mitochondrion |
| P08237 | ATP-dependent 6-phosphofructokinase_ muscle t | Metabolism; Energy pathways | Cytoplasm | 0,000646564 | 2,385281801 | Mitochondrion |
| P09104 | Gamma-enolase | Metabolism; Energy pathways | Cytoplasm | 9,3756E-05 | 0,518397883 | Mitochondrion |
| P09622 | Dihydrolipoyl dehydrogenase_ mitochondrial | Metabolism; Energy pathways | Mitochondrion | 0,016103163 | 1,47064124 | Mitochondrion |
| P0DMV8 | Heat shock 70 kDa protein 1A | Protein Metabolism | Nucleus; Mitochondrion | 0,000412652 | 5,595594618 | Mitochondrion |
| P10515 | Dihydrolipoyllysine-residue acetyltransferase con | Metabolism; Energy pathways | Mitochondrion | 0,029589462 | 2,508495608 | Mitochondrion |
| P10809 | 60 kDa heat shock protein_ mitochondrial | Protein folding | Mitochondrion | 0,044216376 | 0,628073143 | Mitochondrion |
| P11021 | 78 kDa glucose-regulated protein | Protein metabolism | Endoplasmic reticulum; Mitochondrion | 0,000707879 | 2,194011802 | Mitochondrion |
| P11171 | Protein 4.1 | Cell growth | Cytoplasm; Nucleus | 0,001055146 | 2,175784472 | Mitochondrion |
| P11217 | Glycogen phosphorylase_ muscle form | Metabolism; Energy pathways | Endoplasmic reticulum | 0,002388416 | 0,121504671 | Mitochondrion |
| P11586 | C-1-tetrahydrofolate synthase_ cytoplasmic | Metabolism; Energy pathways | Cytoplasm; Mitochondrion | 0,002045822 | 1,910947493 | Mitochondrion |
| P11940 | Polyadenylate-binding protein 1 | Regulation of nucleotide metabolism | Cytoplasm; Nucleus | 0,001859046 | 1,610464782 | Mitochondrion |
| P12235 | ADP/ATP translocase 1 | Transport | Mitochondrion; Nucleus | 0,015863198 | 1,718414432 | Mitochondrion |
| P13010 | X-ray repair cross-complementing protein 5 | Regulation of nucleotide metabolism | Mitochondrion; Nucleus | 0,030430425 | 1,521233982 | Mitochondrion |
| P13645 | Keratin_ type I cytoskeletal 10 | Cell growth | Cytoplasm | 0,040364322 | 0,299399243 | Mitochondrion |
| P13667 | Protein disulfide-isomerase A4 | Protein Metabolism | Endoplasmic reticulum; Mitochondrion | 0,008204204 | 1,983795485 | Mitochondrion |
| P13861 | cAMP-dependent protein kinase type II-alpha reg | Cell communication | Cytoplasm | 0,044099803 | 0,25113101 | Mitochondrion |
| P15880 | 40S ribosomal protein S2 | Protein Metabolism | Ribosome | 0,029035171 | 1,307456401 | Mitochondrion |
| P16401 | Histone H1.5 | Regulation of nucleotide metabolism | Mitochondrion; Nucleus | 0,017619594 | 5,606598823 | Mitochondrion |
| P16435 | NADPH--cytochrome P450 reductase | Metabolism; Energy pathways | Endoplasmic reticulum; Mitochondrion | 0,00672907 | 1,892326594 | Mitochondrion |
| P16615 | Sarcoplasmic/endoplasmic reticulum calcium ATF | Transport | Endoplasmic reticulum; Mitochondrion | 0,004799061 | 2,38561903 | Mitochondrion |
| P17987 | T-complex protein 1 subunit alpha | Protein Metabolism | Cytoplasm | 0,022713229 | 1,439285175 | Mitochondrion |
| P18124 | 60S ribosomal protein L7 | Protein Metabolism | Ribosome | 0,036041691 | 1,229173649 | Mitochondrion |
| P19022 | Cadherin-2 | Cell communication | Plasma membrane | 0,005098477 | 1,900168938 | Mitochondrion |
| P23284 | Peptidyl-prolyl cis-trans isomerase B | Protein Metabolism | Endoplasmic reticulum; Mitochondrion | 0,022412458 | 1,50681539 | Mitochondrion |
| P24539 | ATP synthase F(0) complex subunit B1_ mitochon | Metabolism; Energy pathways | Mitochondrion | 0,045643227 | 1,369621724 | Mitochondrion |
| P25789 | Proteasome subunit alpha type-4 | Protein Metabolism | Cytoplasm; Endoplasmic reticulum | 0,011164883 | 0,596588579 | Mitochondrion |
| P26232 | Catenin alpha-2 | Cell growth | Cytoplasm | 0,001500401 | 1,738755892 | Mitochondrion |
| P26378 | ELAV-like protein 4 | Regulation of nucleotide metabolism | Nucleus | 0,007149181 | 4,321315355 | Mitochondrion |
| P26640 | Valine--tRNA ligase | Protein Metabolism | Cytoplasm; Mitochondrion | 0,020650588 | 1,783375436 | Mitochondrion |
| P26641 | Elongation factor 1-gamma | Protein Metabolism | Cytoplasm; Mitochondrion | 0,016874075 | 0,559968425 | Mitochondrion |
| P27348 | 14-3-3 protein theta | Cell communication | Cytoplasm | 0,006100222 | 0,404830226 | Mitochondrion |
| P27797 | Calreticulin | Protein Metabolism | Endoplasmic reticulum | 0,007835629 | 1,798114751 | Mitochondrion |
| P27816 | Microtubule-associated protein 4 | Cell growth | Cytoplasm | 0,022758961 | 1,760011287 | Mitochondrion |
| P29762 | Cellular retinoic acid-binding protein 1 | Transport | Cytoplasm | 0,016028041 | 0,384281703 | Mitochondrion |
| P30101 | Protein disulfide-isomerase A3 | Protein metabolism | Endoplasmic reticulum | 0,006505876 | 1,702404378 | Mitochondrion |
| P30566 | Adenylosuccinate lyase | Metabolism; Energy pathways | Cytoplasm | 0,004156436 | 0,018234564 | Mitochondrion |
| P31146 | Coronin-1A | Cell growth | Cytoplasm; Mitochondrion | 0,012384327 | 1,59683695 | Mitochondrion |
| P31943 | Heterogeneous nuclear ribonucleoprotein H | Regulation of nucleotide metabolism | Cytoplasm; Nucleus | 0,017331189 | 0,621465691 | Mitochondrion |
| P32969 | 60S ribosomal protein L9 | Protein Metabolism | Ribosome | 0,032647122 | 1,522673981 | Mitochondrion |
| P35908 | Keratin_ type II cytoskeletal 2 epidermal | Cell growth | Cytoplasm | 0,048424202 | 0,289076576 | Mitochondrion |
| P36542 | ATP synthase subunit gamma_ mitochondrial | Metabolism; Energy pathways | Mitochondrion | 0,003299898 | 0,468377323 | Mitochondrion |
| P39656 | Dolichyl-diphosphooligosaccharide--protein glyco | Metabolism; Energy pathways | Endoplasmic reticulum; Mitochondrion | 0,041738795 | 1,389893178 | Mitochondrion |
| P40925 | Malate dehydrogenase_ cytoplasmic | Metabolism; Energy pathways | Cytoplasm; Mitochondrion | 0,002915161 | 0,629553239 | Mitochondrion |
| P40939 | Trifunctional enzyme subunit alpha_ mitochondri | Metabolism; Energy pathways | Mitochondrion | 0,040077104 | 1,60570545 | Mitochondrion |
| P41252 | Isoleucine--tRNA ligase_ cytoplasmic | Protein metabolism | Cytoplasm; Mitochondrion | 0,029308139 | 0,434344756 | Mitochondrion |
| P42766 | 60S ribosomal protein L35 | Protein metabolism | Ribosome | 0,040682623 | 1,502508997 | Mitochondrion |
| P46977 | Dolichyl-diphosphooligosaccharide--protein glycosyltransferase subunit STT3A | | Plasma membrane | 0,008918957 | 2,949041997 | Mitochondrion |
| P48637 | Glutathione synthetase | Metabolism; Energy pathways | Cytoplasm | 0,014073603 | 0,000262862 | Mitochondrion |
| P49327 | Fatty acid synthase | Metabolism; Energy pathways | Cytoplasm; Mitochondrion | 0,040971617 | 0,463236376 | Mitochondrion |
| P49411 | Elongation factor Tu_ mitochondrial | Protein metabolism | Mitochondrion | 0,000903971 | 0,071220585 | Mitochondrion |
| P53007 | Tricarboxylate transport protein_ mitochondrial | Transport | Mitochondrion | 0,004772952 | 1,735069051 | Mitochondrion |
| P54136 | Arginine--tRNA ligase_ cytoplasmic | Protein metabolism | Cytoplasm | 0,048267519 | 1,696283557 | Mitochondrion |
| P54577 | Tyrosine--tRNA ligase_ cytoplasmic | Metabolism; Energy pathways | Cytoplasm | 0,008575453 | 0,44524252 | Mitochondrion |
| P54753 | Ephrin type-B receptor 3 | Cell communication | Plasma membrane | 0,013443429 | 2,179305459 | Mitochondrion |
| P54762 | Ephrin type-B receptor 1 | Cell communication | Plasma membrane | 0,015785663 | 1,673571721 | Mitochondrion |
| P55060 | Exportin-2 | Transport | Nucleus; Mitochondrion | 0,006735833 | 0,077173695 | Mitochondrion |
| P55157 | Microsomal triglyceride transfer protein large sul | Transport | Cytoplasm; Endoplasmic reticulum | 0,009041321 | 3,799179662 | Mitochondrion |
| P59998 | Actin-related protein 2/3 complex subunit 4 | Cell growth | Cytoplasm | 0,040623057 | 0,303683267 | Mitochondrion |
| P60174 | Triosephosphate isomerase | Metabolism; Energy pathways | Cytoplasm | 0,005830156 | 0,579485805 | Mitochondrion |
| P60201 | Myelin proteolipid protein | Cell growth | Plasma membrane | 0,034829337 | 2,010490815 | Mitochondrion |
| P60709 | Actin_ cytoplasmic 1 | Cell growth | Cytoplasm; Mitochondrion | 0,00316421 | 0,279350891 | Mitochondrion |
| P61020 | Ras-related protein Rab-5B | Cell communication | Plasma membrane | 0,015177571 | 0,304565572 | Mitochondrion |
| P61204 | ADP-ribosylation factor 3 | Cell communication | Cytoplasm | 0,007622368 | 0,329778543 | Mitochondrion |
| P61266 | Syntaxin-1B | Transport | Integral to membrane | 0,03150623 | 0,433259888 | Mitochondrion |
| P61313 | 60S ribosomal protein L15 | Protein Metabolism | Ribosome | 0,023308034 | 1,686487303 | Mitochondrion |
| P62140 | Serine/threonine-protein phosphatase PP1-beta c | Cell growth | Nucleus | 0,025656329 | 1,830949891 | Mitochondrion |
| P62424 | 60S ribosomal protein L7a | Protein Metabolism | Cytoplasm; Nucleus | 0,020082076 | 2,746743149 | Mitochondrion |

| | | | | | | |
|---|---|---|---|---|---|---|
| P62495 | Eukaryotic peptide chain release factor subunit 1 | Protein Metabolism | Ribosome | 0,035162562 | 0,11585024 | Mitochondrion |
| P62753 | 40S ribosomal protein S6 | Protein Metabolism | Ribosome | 0,004180199 | 1,482361256 | Mitochondrion |
| P62820 | Ras-related protein Rab-1A | Cell communication | Endoplasmic reticulum; Golgi apparatus | 0,017467777 | 0,416293209 | Mitochondrion |
| P78347 | General transcription factor II-I | Regulation of nucleotide metabolism | Nucleus | 0,04823097 | 2,476103884 | Mitochondrion |
| P78371 | T-complex protein 1 subunit beta | Protein Metabolism | Cytoplasm; Mitochondrion | 0,002477599 | 0,081906731 | Mitochondrion |
| P83731 | 60S ribosomal protein L24 | Protein Metabolism | Ribosome | 0,021746895 | 1,439347593 | Mitochondrion |
| Q00341 | Vigilin | Transport | Cytoplasm | 0,007316173 | 6,463307057 | Mitochondrion |
| Q00534 | Cyclin-dependent kinase 6 | Cell communication | Cytoplasm; Nucleus | 0,019512552 | 2,605122401 | Mitochondrion |
| Q02878 | 60S ribosomal protein L6 | Protein Metabolism | Mitochondrion; Ribosome | 0,00120829 | 1,580017222 | Mitochondrion |
| Q07020 | 60S ribosomal protein L18 | Protein Metabolism | Mitochondrion; Ribosome | 0,015208066 | 1,517185572 | Mitochondrion |
| Q07065 | Cytoskeleton-associated protein 4 | Cell growth | Plasma membrane | 0,024155273 | 1,696189012 | Mitochondrion |
| Q08211 | ATP-dependent RNA helicase A | Regulation of nucleotide metabolism | Cytoplasm; Nucleus | 0,015566351 | 2,020796014 | Mitochondrion |
| Q12926 | ELAV-like protein 2 | Regulation of nucleotide metabolism | Cytoplasm; Nucleus | 0,030745865 | 1,521217526 | Mitochondrion |
| Q13098 | COP9 signalosome complex subunit 1 | Cell communication | Cytoplasm; Nucleus | 0,010461326 | 1,941053905 | Mitochondrion |
| Q13177 | Serine/threonine-protein kinase PAK 2 | Cell communication | Endoplasmic reticulum | 0,036874193 | 1,528754469 | Mitochondrion |
| Q13228 | Selenium-binding protein 1 | Protein Metabolism | Cytoplasm | 0,040256636 | 0,047884205 | Mitochondrion |
| Q13423 | NAD(P) transhydrogenase_ mitochondrial | Metabolism; Energy pathways | Mitochondrion | 0,015457995 | 2,252984583 | Mitochondrion |
| Q13554 | Calcium/calmodulin-dependent protein kinase ty | Cell communication | Cytoplasm | 0,021349714 | 1,582436023 | Mitochondrion |
| Q13557 | Calcium/calmodulin-dependent protein kinase ty | Regulation of cell growth | Cytoplasm; Nucleus | 0,003762239 | 1,785586343 | Mitochondrion |
| Q13838 | Spliceosome RNA helicase DDX39B | Regulation of nucleotide metabolism | Nucleus | 0,031283379 | 0,508874002 | Mitochondrion |
| Q13867 | Bleomycin hydrolase | Metabolism; Energy pathways | Cytoplasm | 0,028741894 | 0,439729631 | Mitochondrion |
| Q14141 | Septin-6 | | Cytoplasm | 0,00686152 | 1,829772259 | Mitochondrion |
| Q14168 | MAGUK p55 subfamily member 2 | Cell communication | Plasma membrane | 0,009262946 | 3,437883514 | Mitochondrion |
| Q14203 | Dynactin subunit 1 | Transport | Cytoplasm; Endoplasmic reticulum | 0,002565822 | 2,383723969 | Mitochondrion |
| Q15084 | Protein disulfide-isomerase A6 | Protein metabolism | Endoplasmic reticulum; Mitochondrion | 0,006800008 | 1,452874978 | Mitochondrion |
| Q15643 | Thyroid receptor-interacting protein 11 | Regulation of nucleotide metabolism | Golgi apparatus | 0,042479717 | 0,503954747 | Mitochondrion |
| Q16537 | Serine/threonine-protein phosphatase 2A 56 kDa | Cell communication | Cytoplasm | 0,001348344 | 2,227729546 | Mitochondrion |
| Q5ZPR3 | CD276 antigen | | | 0,007724194 | 0,568710857 | Mitochondrion |
| Q6P2E9 | Enhancer of mRNA-decapping protein 4 | | Nucleus | 0,047087947 | 0,275246914 | Mitochondrion |
| Q6S8J3 | POTE ankyrin domain family member E | | Plasma membrane | 0,000335496 | 0,289606564 | Mitochondrion |
| Q6XQN6 | Nicotinate phosphoribosyltransferase | Regulation of nucleotide metabolism | Cytoplasm | 0,026969894 | 0,115305526 | Mitochondrion |
| Q71U36 | Tubulin alpha-1A chain | Cell growth | Cytoplasm; Nucleus | 0,016442503 | 0,391012742 | Mitochondrion |
| Q8N3J6 | Cell adhesion molecule 2 | Cell communication | | 4,80216E-05 | 2,178052746 | Mitochondrion |
| Q8N9N7 | Leucine-rich repeat-containing protein 57 | | | 0,001793816 | 1,504067232 | Mitochondrion |
| Q8NC51 | Plasminogen activator inhibitor 1 RNA-binding pr | Regulation of nucleotide metabolism | Cytoplasm | 0,015575693 | 2,180715454 | Mitochondrion |
| Q8NDA2 | Hemicentin-2 | | | 0,031894072 | 1,798072069 | Mitochondrion |
| Q92499 | ATP-dependent RNA helicase DDX1 | Regulation of nucleotide metabolism | Nucleus | 0,015459682 | 1,841998691 | Mitochondrion |
| Q92973 | Transportin-1 | Transport | Cytoplasm | 0,014955752 | 1,801288606 | Mitochondrion |
| Q96CS3 | FAS-associated factor 2 | | Cytoplasm | 0,010165185 | 2,158472303 | Mitochondrion |
| Q96CW1 | AP-2 complex subunit mu | Transport | | 0,038049452 | 4,082798388 | Mitochondrion |
| Q96E17 | Ras-related protein Rab-3C | Transport | Cytoplasm | 0,001939255 | 2,227435164 | Mitochondrion |
| Q99798 | Aconitate hydratase_ mitochondrial | Metabolism; Energy pathways | Mitochondrion | 0,02269079 | 2,215956422 | Mitochondrion |
| Q9BQE3 | Tubulin alpha-1C chain | Cell growth | Cytoplasm; Nucleus | 0,000122696 | 0,598538737 | Mitochondrion |
| Q9BUF5 | Tubulin beta-6 chain | Cell growth | | 0,021991092 | 0,124754562 | Mitochondrion |
| Q9BZF1 | Oxysterol-binding protein-related protein 8 | Transport | Mitochondrion | 0,001967672 | 0,284492675 | Mitochondrion |
| Q9C0E8 | Endoplasmic reticulum junction formation protein lunapark | | | 0,002615647 | 8,365588447 | Mitochondrion |
| Q9H115 | Beta-soluble NSF attachment protein | Transport | | 0,009873899 | 3,517510782 | Mitochondrion |
| Q9H270 | Vacuolar protein sorting-associated protein 11 ho | Transport | Endosome | 0,047816398 | 0,581616808 | Mitochondrion |
| Q9HDC9 | Adipocyte plasma membrane-associated protein | | Plasma membrane | 0,024978764 | 2,203807825 | Mitochondrion |
| Q9NQC3 | Reticulon-4 | Cell growth | Endoplasmic reticulum; Mitochondrion | 0,013811803 | 1,360480198 | Mitochondrion |
| Q9NSD9 | Phenylalanine--tRNA ligase beta subunit | Protein Metabolism | Cytoplasm | 0,039754019 | 2,109187107 | Mitochondrion |
| Q9NTJ5 | Phosphatidylinositide phosphatase SAC1 | Cell communication | Endoplasmic reticulum | 0,028133814 | 4,765291663 | Mitochondrion |
| Q9NVA2 | Septin-11 | Cell cycle | Cytoplasm | 0,000575627 | 2,377740197 | Mitochondrion |
| Q9NYU2 | UDP-glucose:glycoprotein glucosyltransferase 1 | Metabolism; Energy pathways | Endoplasmic reticulum; Mitochondrion | 0,004860355 | 0,33513755 | Mitochondrion |
| Q9NZI8 | Insulin-like growth factor 2 mRNA-binding protein 1 | | | 0,000457846 | 1,857132891 | Mitochondrion |
| Q9P121 | Neurotrimin | | | 0,000222943 | 0,595228229 | Mitochondrion |
| Q9UHD8 | Septin-9 | | | 0,004379017 | 1,837788151 | Mitochondrion |
| Q9UHG3 | Prenylcysteine oxidase 1 | | | 0,022543367 | 1,707597825 | Mitochondrion |
| Q9UI15 | Transgelin-3 | Regulation of transcription | Cytoplasm; Nucleus | 0,017132729 | 1,650897189 | Mitochondrion |
| Q9UIW2 | Plexin-A1 | Cell communication | Plasma membrane | 0,009879354 | 0,144275572 | Mitochondrion |
| Q9UKA9 | Polypyrimidine tract-binding protein 2 | | | 0,048598456 | 2,387963071 | Mitochondrion |
| Q9UL18 | Protein argonaute-1 | Protein Metabolism | Cytoplasm | 0,017594549 | 3,572888237 | Mitochondrion |
| Q9ULU8 | Calcium-dependent secretion activator 1 | Transport | Cytoplasm | 5,14431E-08 | 3,651620072 | Mitochondrion |
| Q9UM54 | Unconventional myosin-VI | Cell growth | Golgi apparatus | 0,030425256 | 2,38182267 | Mitochondrion |
| Q9UPZ6 | Thrombospondin type-1 domain-containing prote | Cell communication | | 0,000246963 | 0,436256441 | Mitochondrion |
| Q9UQ80 | Proliferation-associated protein 2G4 | Regulation of nucleotide metabolism | Mitochondrion; Nucleus | 0,009869115 | 0,532020767 | Mitochondrion |
| Q9Y265 | RuvB-like 1 | Regulation of nucleotide metabolism | Cytoplasm; Nucleus | 0,016113411 | 0,148128232 | Mitochondrion |
| Q9Y2A7 | Nck-associated protein 1 | Cell communication | Cytoplasm; Mitochondrion | 0,02506563 | 0,695673509 | Mitochondrion |
| Q9Y4F1 | FERM_ ARHGEF and pleckstrin domain-containing | Cell growth | | 0,002056227 | 2,34627575 | Mitochondrion |
| Q9Y6G9 | Cytoplasmic dynein 1 light intermediate chain 1 | Cell growth | Cytoplasm | 0,021376249 | 1,425989929 | Mitochondrion |
| Q9Y6M1 | Insulin-like growth factor 2 mRNA-binding protei | Regulation of nucleotide metabolism | Cytoplasm | 0,000884164 | 2,245365845 | Mitochondrion |
| O00231 | 26S proteasome non-ATPase regulatory subunit 1 | Protein Metabolism | Cytoplasm; Nucleus | 0,042674718 | 0,759190435 | Nucleus |
| O00232 | 26S proteasome non-ATPase regulatory subunit 1 | Protein Metabolism | Cytoplasm | 0,006864619 | 0,646012592 | Nucleus |
| O00299 | Chloride intracellular channel protein 1 | Transport | Nucleus | 0,008515495 | 0,453071903 | Nucleus |
| O43175 | D-3-phosphoglycerate dehydrogenase | Metabolism; Energy pathways | Extracellular | 0,042751498 | 0,633175248 | Nucleus |
| O43488 | Aflatoxin B1 aldehyde reductase member 2 | Metabolism; Energy pathways | Cytoplasm | 0,012044747 | 0,7203363 | Nucleus |
| O43707 | Alpha-actinin-4 | Cell growth | Cytoplasm; Nucleus | 0,047269207 | 3,277134376 | Nucleus |
| O43776 | Asparagine--tRNA ligase_ cytoplasmic | Protein Metabolism | Cytoplasm | 0,011291814 | 0,55322836 | Nucleus |
| O43854 | EGF-like repeat and discoidin I-like domain-contai | Cell growth | Extracellular | 0,046525395 | 0,098187089 | Nucleus |
| O60282 | Kinesin heavy chain isoform 5C | Cell growth | Cytoplasm | 0,023907322 | 2,437747158 | Nucleus |
| O60701 | UDP-glucose 6-dehydrogenase | Metabolism; Energy pathways | Nucleus | 0,005454333 | 1,952782372 | Nucleus |
| O60812 | Heterogeneous nuclear ribonucleoprotein C-like 1 | Protein Metabolism | Nucleus | 0,001446525 | 1,247542424 | Nucleus |
| O75122 | CLIP-associating protein 2 | Cell growth | Cytoplasm | 0,001610965 | 2,963054274 | Nucleus |
| O75746 | Calcium-binding mitochondrial carrier protein Ara | Transport | Cytoplasm; Mitochondrion | 0,031292387 | 1,956705671 | Nucleus |
| O94973 | AP-2 complex subunit alpha-2 | Transport | Plasma membrane | 0,04772019 | 1,694377982 | Nucleus |
| O94986 | Centrosomal protein of 152 kDa | Cell growth | Cytoplasm; Nucleus | 0,038838029 | 2,376862093 | Nucleus |
| P00568 | Adenylate kinase isoenzyme 1 | Metabolism; Energy pathways | Cytoplasm | 0,0312693 | 0,50159433 | Nucleus |
| P04040 | Catalase | Metabolism; Energy pathways | Cytoplasm | 0,01498074 | 0,309666879 | Nucleus |
| P05023 | Sodium/potassium-transporting ATPase subunit a | Transport | Plasma membrane | 0,042951435 | 1,759581853 | Nucleus |
| P06744 | Glucose-6-phosphate isomerase | Metabolism; Energy pathways | Cytoplasm | 0,00151362 | 2,739221368 | Nucleus |
| P07305 | Histone H1.0 | Regulation of nucleotide metabolism | Nucleus | 0,01003865 | 0,17174482 | Nucleus |
| P07339 | Cathepsin D | Protein Metabolism | Lysosome | 0,00971599 | 1,453197247 | Nucleus |
| P07355 | Putative annexin A2-like protein | Signal transduction; Cell communication | Nucleus | 0,001676842 | 0,117188794 | Nucleus |
| P07437 | Tubulin beta chain | Cell growth | Cytoplasm; Plasma membrane | 0,034092144 | 0,562841615 | Nucleus |
| P08865 | 40S ribosomal protein SA | Protein Metabolism | Nucleus | 0,028549503 | 0,537863359 | Nucleus |
| P09471 | Guanine nucleotide-binding protein G(o) subunit | Cell communication | Plasma membrane | 6,63892E-05 | 2,281275734 | Nucleus |
| P10909 | Clusterin | Protein metabolism | Cytoplasm; Nucleus | 0,012455714 | 2,501360621 | Nucleus |
| P11940 | Polyadenylate-binding protein 1 | Regulation of nucleotide metabolism | Cytoplasm; Nucleus | 0,007646054 | 0,751427291 | Nucleus |
| P12004 | Proliferating cell nuclear antigen | DNA repair | Nucleus | 0,012781361 | 0,392478613 | Nucleus |

| | | | | | | |
|---|---|---|---|---|---|---|
| P12235 | ADP/ATP translocase 1 | Transport | Mitochondrion; Nucleus | 0,022048751 | 1,330283404 | Nucleus |
| P12236 | ADP/ATP translocase 3 | Transport | Mitochondrion; Nucleus | 0,031299549 | 1,263580596 | Nucleus |
| P13667 | Protein disulfide-isomerase A4 | Protein Metabolism | Endoplasmic reticulum; Mitochondrion | 0,009831539 | 1,831403121 | Nucleus |
| P14314 | Glucosidase 2 subunit beta | Metabolism; Energy pathways | Endoplasmic reticulum | 0,005705199 | 2,070427651 | Nucleus |
| P15311 | Ezrin | Cell growth | Cytoplasm | 0,019489594 | 1,64373973 | Nucleus |
| P15586 | N-acetylglucosamine-6-sulfatase | Metabolism; Energy pathways | Lysosome | 0,008373919 | 1,943994832 | Nucleus |
| P18621 | 60S ribosomal protein L17 | Protein Metabolism | Ribosome | 0,033544525 | 0,220084069 | Nucleus |
| P19022 | Cadherin-2 | Cell communication | Plasma membrane | 0,002881189 | 2,752353198 | Nucleus |
| P19367 | Hexokinase-1 | Metabolism; Energy pathways | Cytoplasm | 0,03430051 | 1,971098315 | Nucleus |
| P21579 | Synaptotagmin-1 | Cell communication | Cytoplasm | 0,012292234 | 2,422322432 | Nucleus |
| P23526 | Adenosylhomocysteinase | Metabolism; Energy pathways | Cytoplasm | 0,035066789 | 0,803159232 | Nucleus |
| P26232 | Catenin alpha-2 | Cell growth | Cytoplasm | 0,016305754 | 1,569844886 | Nucleus |
| P26378 | ELAV-like protein 4 | Regulation of nucleotide metabolism | Nucleus | 0,007170079 | 1,856229718 | Nucleus |
| P26640 | Valine--tRNA ligase | Protein Metabolism | Cytoplasm; Mitochondrion | 0,046815094 | 3,808227234 | Nucleus |
| P27824 | Calnexin | Protein folding | Endoplasmic reticulum | 0,042193938 | 1,633185619 | Nucleus |
| P29323 | Ephrin type-B receptor 2 | Cell communication | Plasma membrane | 0,027360125 | 1,320946438 | Nucleus |
| P29762 | Cellular retinoic acid-binding protein 1 | Transport | Cytoplasm | 0,012305218 | 0,436782176 | Nucleus |
| P31150 | Rab GDP dissociation inhibitor alpha | Cell communication | Cytoplasm | 0,005300074 | 0,611715363 | Nucleus |
| P31939 | Bifunctional purine biosynthesis protein PURH | Metabolism; Energy pathways | Cytoplasm; Mitochondrion | 0,001149528 | 0,724149889 | Nucleus |
| P32969 | 60S ribosomal protein L9 | Protein Metabolism | Ribosome | 0,018378246 | 0,688091323 | Nucleus |
| P35221 | Catenin alpha-1 | Cell growth | Cytoplasm | 0,027020224 | 2,07107355 | Nucleus |
| P35222 | Catenin beta-1 | Cell communication | Nucleus; Plasma membrane | 0,008298486 | 3,084534228 | Nucleus |
| P35611 | Alpha-adducin | Cell growth | Nucleus | 0,003428561 | 2,265544866 | Nucleus |
| P35612 | Beta-adducin | Cell growth | Nucleus | 0,004047981 | 2,744406909 | Nucleus |
| P36578 | 60S ribosomal protein L4 | Protein Metabolism | Nucleus; Ribosome | 0,03625426 | 0,657697261 | Nucleus |
| P36776 | Lon protease homolog_ mitochondrial | Protein Metabolism | Mitochondrion | 0,013581566 | 2,149800119 | Nucleus |
| P38159 | RNA-binding motif protein_ X chromosome | Regulation of nucleotide metabolism | Nucleus | 0,044884793 | 1,286154056 | Nucleus |
| P41091 | Putative eukaryotic translation initiation factor 2 | Protein Metabolism | Cytoplasm; Nucleus | 0,045071455 | 1,610767543 | Nucleus |
| P45880 | Voltage-dependent anion-selective channel prote | Transport | Mitochondrion; Nucleus | 0,02356102 | 1,272432256 | Nucleus |
| P46776 | 60S ribosomal protein L27a | Protein Metabolism | Nucleus; Ribosome | 0,013756968 | 0,518782338 | Nucleus |
| P46781 | 40S ribosomal protein S9 | Protein Metabolism | Nucleus; Ribosome | 0,004285354 | 0,674963642 | Nucleus |
| P46783 | 40S ribosomal protein S10 | Protein Metabolism | Nucleus; Ribosome | 0,033328552 | 0,432343805 | Nucleus |
| P48681 | Nestin | Protein Metabolism | Cytoplasm; Nucleus | 0,001442497 | 0,514569526 | Nucleus |
| P49591 | Serine--tRNA ligase_ cytoplasmic | Metabolism; Energy pathways | Cytoplasm | 0,007789969 | 0,742013578 | Nucleus |
| P49841 | Glycogen synthase kinase-3 beta | Metabolism; Energy pathways | Cytoplasm; Nucleus | 0,028793336 | 1,923916175 | Nucleus |
| P50395 | Rab GDP dissociation inhibitor beta | Transport | Cytoplasm | 0,032959278 | 1,196666575 | Nucleus |
| P50454 | Serpin H1 | Protein Metabolism | Endoplasmic reticulum; Nucleus | 0,001882583 | 0,273963977 | Nucleus |
| P50914 | 60S ribosomal protein L14 | Protein Metabolism | Nucleus; Ribosome | 0,01397447 | 0,664602733 | Nucleus |
| P51991 | Heterogeneous nuclear ribonucleoprotein A3 | Regulation of nucleotide metabolism | Nucleus | 0,03162002 | 0,241198256 | Nucleus |
| P52209 | 6-phosphogluconate dehydrogenase_ decarboxyl | Metabolism; Energy pathways | Cytoplasm | 0,02555359 | 0,642189831 | Nucleus |
| P52292 | Importin subunit alpha-1 | Cell communication | Cytoplasm; Nucleus | 0,011475161 | 0,53527724 | Nucleus |
| P52895 | Aldo-keto reductase family 1 member C2 | Metabolism; Transport | Cytoplasm | 0,000233538 | 1,652717252 | Nucleus |
| P54578 | Ubiquitin carboxyl-terminal hydrolase 14 | Protein Metabolism | Cytoplasm | 0,036102138 | 0,793014278 | Nucleus |
| P54687 | Branched-chain-amino-acid aminotransferase_ cy | Metabolism; Energy pathways | Cytoplasm | 0,026474716 | 0,63455212 | Nucleus |
| P54920 | Alpha-soluble NSF attachment protein | Transport | Golgi apparatus | 0,041703561 | 1,250832907 | Nucleus |
| P55265 | Double-stranded RNA-specific adenosine deamin | Regulation of nucleotide metabolism | Nucleus | 0,030038645 | 2,229249735 | Nucleus |
| P55809 | Succinyl-CoA:3-ketoacid coenzyme A transferase | Metabolism; Energy pathways | Mitochondrion; Nucleus | 0,003366947 | 2,312218782 | Nucleus |
| P60953 | Cell division control protein 42 homolog | Cell communication | Cytoplasm | 0,019566354 | 1,234811874 | Nucleus |
| P61247 | 40S ribosomal protein S3a | Protein Metabolism | Cytoplasm; Nucleus | 0,007898504 | 0,587101564 | Nucleus |
| P61313 | 60S ribosomal protein L15 | Protein Metabolism | Ribosome | 0,003285851 | 0,641397739 | Nucleus |
| P61421 | V-type proton ATPase subunit d 1 | Transport | | 0,011668732 | 1,359521955 | Nucleus |
| P62241 | 40S ribosomal protein S8 | Protein Metabolism | Cytoplasm; Nucleus | 0,008648868 | 0,607418512 | Nucleus |
| P62263 | 40S ribosomal protein S14 | Protein Metabolism | Nucleus; Ribosome | 0,00155768 | 0,373603862 | Nucleus |
| P62277 | 40S ribosomal protein S13 | Protein Metabolism | Nucleus; Ribosome | 0,032199766 | 0,464835475 | Nucleus |
| P62280 | 40S ribosomal protein S11 | Protein Metabolism | Nucleus; Ribosome | 0,000594412 | 0,525262296 | Nucleus |
| P62424 | 60S ribosomal protein L7a | Protein Metabolism | Cytoplasm; Nucleus | 0,028859607 | 0,606764491 | Nucleus |
| P62701 | 40S ribosomal protein S4_ X isoform | Protein Metabolism | Nucleus; Ribosome | 0,032482507 | 0,599988502 | Nucleus |
| P62736 | Actin_ aortic smooth muscle | Cell growth | Cytoplasm | 0,019775708 | 0,599234996 | Nucleus |
| P62826 | GTP-binding nuclear protein Ran | Cell communication | Nucleus | 0,013095323 | 0,698407722 | Nucleus |
| P62879 | Guanine nucleotide-binding protein G(I)/G(S)/G(T | Cell communication | Cytoplasm | 0,037159381 | 1,508968061 | Nucleus |
| P62910 | 60S ribosomal protein L32 | Protein Metabolism | Ribosome | 0,018938121 | 0,707083307 | Nucleus |
| P62913 | 60S ribosomal protein L11 | Protein Metabolism | Nucleus; Ribosome | 0,011005093 | 0,709373454 | Nucleus |
| P62987 | Ubiquitin-60S ribosomal protein L40 | Protein Metabolism | Ribosome | 0,02900742 | 1,518431693 | Nucleus |
| P63244 | Receptor of activated protein C kinase 1 | Cell communication | Cytoplasm; Nucleus | 0,044191724 | 0,718381678 | Nucleus |
| P68402 | Platelet-activating factor acetylhydrolase IB subu | Cell communication | Cytoplasm | 0,005227115 | 0,077812948 | Nucleus |
| Q00169 | Phosphatidylinositol transfer protein alpha isofor | Transport | Cytoplasm | 0,00054813 | 0,649047536 | Nucleus |
| Q01518 | Adenylyl cyclase-associated protein 1 | Cell growth | Cytoplasm; Nucleus | 0,002032923 | 1,672220377 | Nucleus |
| Q02790 | Peptidyl-prolyl cis-trans isomerase FKBP4 | Metabolism; Energy pathways | Cytoplasm; Nucleus | 0,019691596 | 1,549846489 | Nucleus |
| Q02978 | Mitochondrial 2-oxoglutarate/malate carrier prot | Transport | Mitochondrion; Nucleus | 0,018543913 | 1,596987798 | Nucleus |
| Q07955 | Serine/arginine-rich splicing factor 1 | Protein Metabolism | Nucleus | 0,006410167 | 0,011706407 | Nucleus |
| Q08209 | Serine/threonine-protein phosphatase 2B catalyti | Cell communication | Cytoplasm; Nucleus | 0,005333659 | 1,674281379 | Nucleus |
| Q10567 | AP-1 complex subunit beta-1 | Transport | Golgi apparatus | 0,049241385 | 1,577211384 | Nucleus |
| Q12906 | Interleukin enhancer-binding factor 3 | Regulation of gene expression | Nucleus | 0,0416942 | 1,476921817 | Nucleus |
| Q13098 | COP9 signalosome complex subunit 1 | Cell communication | Cytoplasm; Nucleus | 0,00301648 | 0,460558086 | Nucleus |
| Q14108 | Lysosome membrane protein 2 | Cell communication | Plasma membrane | 0,012894874 | 2,224613645 | Nucleus |
| Q14117 | Dihydropyrimidinase | Regulation of nucleotide metabolism | Cytoplasm; Nucleus | 0,031118586 | 1,770374745 | Nucleus |
| Q14204 | Cytoplasmic dynein 1 heavy chain 1 | Metabolism; Energy pathways | Cytoplasm; Nucleus | 0,00646492 | 0,822854117 | Nucleus |
| Q15102 | Platelet-activating factor acetylhydrolase IB subu | Metabolism; Energy pathways | Cytoplasm | 0,039944965 | 0,366582625 | Nucleus |
| Q15185 | Prostaglandin E synthase 3 | Protein Metabolism | Cytoplasm; Nucleus | 0,00429819 | 0,643450664 | Nucleus |
| Q15334 | Lethal(2) giant larvae protein homolog 1 | Cell growth | Cytoplasm | 0,030107933 | 2,281544421 | Nucleus |
| Q16531 | DNA damage-binding protein 1 | Regulation of nucleotide metabolism | Nucleus | 0,008915944 | 0,299821511 | Nucleus |
| Q5TF21 | Protein SOGA3 | | Plasma membrane | 0,032542655 | 2,382704942 | Nucleus |
| Q5VTE0 | Putative elongation factor 1-alpha-like 3 | | | 0,00620493 | 0,801519443 | Nucleus |
| Q71U36 | Tubulin alpha-1A chain | Cell growth | Cytoplasm; Nucleus | 0,042963242 | 0,720965363 | Nucleus |
| Q7L099 | Protein RUFY3 | | | 0,030921726 | 1,975790883 | Nucleus |
| Q7Z460 | CLIP-associating protein 1 | Cell growth | Nucleus | 0,024255639 | 1,959291554 | Nucleus |
| Q86UX7 | Fermitin family homolog 3 | | Plasma membrane | 0,035330665 | 2,750294859 | Nucleus |
| Q8N163 | Cell cycle and apoptosis regulator protein 2 | Signal transduction | Nucleus | 0,01716188 | 0,656241816 | Nucleus |
| Q8WVM8 | Sec1 family domain-containing protein 1 | Transport | Nucleus; Plasma membrane | 0,010468417 | 1,740028619 | Nucleus |
| Q8WZA9 | Immunity-related GTPase family Q protein | | | 0,043640359 | 1,488348652 | Nucleus |
| Q92820 | Gamma-glutamyl hydrolase | Metabolism; Energy pathways | Lysosome | 2,13731E-05 | 3,403527114 | Nucleus |
| Q96JP2 | Unconventional myosin-XVB | | | 0,000665393 | 2,247191517 | Nucleus |
| Q96KP4 | Cytosolic non-specific dipeptidase | Protein Metabolism | Cytoplasm | 0,018082124 | 0,653609793 | Nucleus |
| Q96QT4 | Transient receptor potential cation channel subfa | Transport | Plasma membrane | 0,048510167 | 1,772793096 | Nucleus |
| Q99460 | 26S proteasome non-ATPase regulatory subunit 1 | Protein Metabolism | Cytoplasm; Nucleus | 0,000178441 | 0,534552509 | Nucleus |
| Q99497 | Protein/nucleic acid deglycase DJ-1 | Regulation of nucleotide metabolism | Nucleus | 0,035666701 | 1,842977904 | Nucleus |
| Q99873 | Protein arginine N-methyltransferase 1 | Metabolism; Energy pathways | Nucleus | 0,049939145 | 0,646011394 | Nucleus |
| Q99996 | A-kinase anchor protein 9 | Cell communication | Cytoplasm | 8,85179E-07 | 2,036444024 | Nucleus |
| Q9BSJ8 | Extended synaptotagmin-1 | | | 0,02619189 | 0,614668628 | Nucleus |

| | | | | | | |
|---|---|---|---|---|---|---|
| Q9BVA1 | Tubulin beta-2B chain | Cell growth | Cytoplasm; Nucleus | 0,037178297 | 0,544267532 | Nucleus |
| Q9H9A6 | Leucine-rich repeat-containing protein 40 | | | 0,029212075 | 1,840183043 | Nucleus |
| Q9H9B4 | Sideroflexin-1 | Transport | Mitochondrion | 0,030568829 | 1,288703449 | Nucleus |
| Q9HDC9 | Adipocyte plasma membrane-associated protein | | | 0,007445066 | 0,822975776 | Nucleus |
| Q9NR30 | Nucleolar RNA helicase 2 | Transcription | Nucleus | 0,042639097 | 2,016705175 | Nucleus |
| Q9NRF8 | CTP synthase 2 | Regulation of nucleotide metabolism | Nucleus | 0,004050885 | 0,6756135 | Nucleus |
| Q9NTK5 | Obg-like ATPase 1 | | Nucleus | 0,001003329 | 0,540163112 | Nucleus |
| Q9NVA2 | Septin-11 | Cell cycle | Cytoplasm | 0,029013932 | 1,526452672 | Nucleus |
| Q9UGL1 | Lysine-specific demethylase 5B | Regulation of nucleotide metabolism | Nucleus | 0,041021724 | 2,003951507 | Nucleus |
| Q9UI15 | Transgelin-3 | Regulation of transcription | Cytoplasm; Nucleus | 0,011143049 | 1,826828779 | Nucleus |
| Q9UJS0 | Calcium-binding mitochondrial carrier protein Ara | Transport | Mitochondrion | 0,009323279 | 1,509964493 | Nucleus |
| Q9UKX3 | Myosin-13 | Cell growth | Cytoplasm | 0,026295856 | 0,561942769 | Nucleus |
| Q9UQE7 | Structural maintenance of chromosomes protein | DNA repair | Nucleus | 0,038980862 | 1,73005412 | Nucleus |
| Q9Y277 | Voltage-dependent anion-selective channel prote | Transport | Mitochondrion | 0,003551196 | 1,255875827 | Nucleus |
| Q9Y4F4 | TOG array regulator of axonemal microtubules protein 1 | | | 0,016199351 | 2,085827533 | Nucleus |
| Q9Y617 | Phosphoserine aminotransferase | Metabolism; Energy pathways | Cytoplasm | 7,79348E-05 | 0,482136366 | Nucleus |
| Q9Y6C9 | Mitochondrial carrier homolog 2 | Cell communication | Mitochondrion | 0,007181508 | 1,545978896 | Nucleus |
