## Supplementary Table 3 for "Mitochondrial, cell cycle control and neuritogenesis alterations in an iPSC-based neurodevelopmental model for schizophrenia"

| Accession | Description | Biological Process | Cellular component | p-Value | SCZ/CTRL ration |
|---|---|---|---|---|---|
| A4FU69 | EF-hand calcium-binding domain-containing protein 5 | | | 0,001101 | 2,724427411 |
| A4UGR9 | Xin actin-binding repeat-containing protein 2 | Cytoskeletal anchoring | Cytoplasm | 0,006756 | 1,413953388 |
| A6NCE7 | Microtubule-associated proteins 1A/1B light chain 3 beta 2 | | Cytoplasm | 0,001417 | 1,99612969 |
| B2RPK0 | Putative high mobility group protein B1-like 1 | | | 0,032314 | 1,590743352 |
| O00231 | 26S proteasome non-ATPase regulatory subunit 11 | Protein metabolism | Cytoplasm; Nucleus | 0,000029 | 1,428664042 |
| O00232 | 26S proteasome non-ATPase regulatory subunit 12 | Protein metabolism | Cytoplasm | 0,008566 | 1,544545922 |
| O00264 | Membrane-associated progesterone receptor component 1 | Cell communication | Plasma membrane | 0,001459 | 2,322924147 |
| O00429 | Dynamin-1-like protein | Mitochondrion organization and biogenesis | Cytoplasm | 0,006560 | 0,06391487 |
| O00567 | Nucleolar protein 56 | Regulation of nucleotide metabolism | Nucleus | 0,007330 | 3,842896338 |
| O14737 | Programmed cell death protein 5 | Apoptosis | Cytoplasm; Nucleus | 0,006358 | 3,836727237 |
| O14818 | Proteasome subunit alpha type-7 | Protein metabolism | Cytoplasm | 0,030521 | 1,893928387 |
| O14979 | Heterogeneous nuclear ribonucleoprotein D-like | Regulation of nucleotide metabolism | Nucleus | 0,000637 | 2,150005885 |
| O15078 | Centrosomal protein of 290 kDa | | | 0,015359 | 14,26648619 |
| O15347 | High mobility group protein B3 | Regulation of nucleotide metabolism | Nucleus | 0,005500 | 1,364309014 |
| O15540 | Fatty acid-binding protein_ brain | Transport | Cytoplasm | 0,000087 | 3,125786118 |
| O43237 | Cytoplasmic dynein 1 light intermediate chain 2 | | Cytoplasm | 0,000001 | 2,899117004 |
| O43390 | Heterogeneous nuclear ribonucleoprotein R | Regulation of nucleotide metabolism | Nucleus | 0,008258 | 3,137959143 |
| O43765 | Small glutamine-rich tetratricopeptide repeat-containing protein alpha | Protein metabolism | Cytoplasm | 0,032581 | 1,576418524 |
| O60282 | Kinesin heavy chain isoform 5C | Transport | Cytoeskeleton | 0,000039 | 1,423144029 |
| O60506 | Heterogeneous nuclear ribonucleoprotein Q | Regulation of nucleotide metabolism | Cytoplasm | 0,003433 | 1,530731557 |
| O75131 | Copine-3 | Transport | Cytoplasm | 0,000171 | 3,456872861 |
| O75347 | Tubulin-specific chaperone A | Protein metabolism | Cytoskeleton | 0,000026 | 3,840566927 |
| O75382 | Tripartite motif-containing protein 3 | Cell growth | Cytoplasm | 0,026096 | 0,076976149 |
| O75390 | Citrate synthase_ mitochondrial | Metabolism; Energy pathways | Cytoplasm; Mitochondrion | 0,004420 | 2,646990187 |
| O75396 | Vesicle-trafficking protein SEC22b | Cell communication | Endoplasmic reticulum; Mitoch | 0,012905 | 0,683726194 |
| O75475 | PC4 and SFRS1-interacting protein | Regulation of nucleotide metabolism | Cytoplasm; Nucleus | 0,005454 | 1,833569047 |
| O75874 | Isocitrate dehydrogenase [NADP] cytoplasmic | Metabolism; Energy pathways | Cytoplasm | 0,002529 | 1,683910315 |
| O94986 | Centrosomal protein of 152 kDa | | Centrosome | 0,031185 | 1,692258676 |
| O95084 | Serine protease 23 | Protein metabolism | | 0,017703 | 0,317260583 |
| O95232 | Luc7-like protein 3 | Regulation of nucleotide metabolism | Nucleus | 0,018226 | 0,514816495 |
| O95336 | 6-phosphogluconolactonase | Metabolism; Energy pathways | | 0,046459 | 5,101197384 |
| O95747 | Serine/threonine-protein kinase R1 | Cell communication | Cytoplasm | 0,000378 | 1,525816307 |
| O95865 | N(G)_N(G)-dimethylarginine dimethylaminohydrolase 2 | Metabolism; Energy pathways | Cytoplasm | 0,000182 | 3,075603829 |
| O96006 | Zinc finger BED domain-containing protein 1 | Regulation of nucleotide metabolism | Nucleus | 0,007524 | 2,052376373 |
| P00338 | L-lactate dehydrogenase A chain | Metabolism; Energy pathways | Cytoplasm | 0,006554 | 1,683082038 |
| P00367 | Glutamate dehydrogenase 1_ mitochondrial | Metabolism; Energy pathways | Mitochondrion | 0,047704 | 1,352938725 |
| P00441 | Superoxide dismutase [Cu-Zn] | Metabolism; Energy pathways | Cytoplasm | 0,001274 | 5,280017128 |
| P00558 | Phosphoglycerate kinase 1 | Metabolism; Energy pathways | Cytoplasm | 0,021835 | 2,656123884 |
| P00568 | Adenylate kinase isoenzyme 1 | Metabolism; Energy pathways | Cytoplasm | 0,008022 | 1,675044517 |
| P02768 | Serum albumin | Transport | Extracellular | 0,008951 | 2,143526733 |
| P04075 | Fructose-bisphosphate aldolase A | Metabolism; Energy pathways | Cytoplasm | 0,033034 | 1,533279394 |
| P04350 | Tubulin beta-4A chain | Cell growth | Microtubule | 0,039634 | 1,451155651 |
| P04406 | Glyceraldehyde-3-phosphate dehydrogenase | Metabolism; Energy pathways | Cytoplasm | 0,012201 | 1,8649441 |
| P05387 | 60S acidic ribosomal protein P2 | Protein metabolism | Cytoplasm | 0,000310 | 4,347939219 |
| P05937 | Calbindin | Cell communication | Cytoplasm; Nucleus | 0,000001 | 5,206278496 |
| P06733 | Alpha-enolase | Metabolism; Energy pathways | Cytoplasm | 0,006600 | 2,257962433 |
| P06744 | Glucose-6-phosphate isomerase | Metabolism; Energy pathways | Cytoplasm | 0,008231 | 1,914536761 |
| P06753 | Tropomyosin alpha-3 chain | Cell growth | Cytoplasm | 0,002648 | 1,990681561 |
| P07195 | L-lactate dehydrogenase B chain | Metabolism; Energy pathways | Cytoplasm | 0,004090 | 1,713553818 |
| P07305 | Histone H1.0 | Regulation of nucleotide metabolism | Nucleus | 0,009729 | 0,194459798 |
| P07339 | Cathepsin D | Protein metabolism | Lysosome | 0,004048 | 1,851891243 |
| P07437 | Tubulin beta chain | Cell growth | Cytoplasm | 0,004369 | 1,818524126 |
| P07602 | Prosaposin | Cell communication | Lysosome | 0,000001 | 2,641685059 |
| P07737 | Profilin-1 | Cell growth | Cytoplasm | 0,019891 | 2,656841125 |
| P07864 | L-lactate dehydrogenase C chain | Metabolism; Energy pathways | Cytoplasm | 0,000025 | 1,825307604 |
| P08133 | Annexin A6 | Cell communication | Endoplasmic reticulum | 0,021518 | 1,900748397 |
| P08727 | Keratin_ type I cytoskeletal 19 | Cell growth | Cytoplasm | 0,046869 | 0,65532044 |
| P09104 | Gamma-enolase | Metabolism; Energy pathways | Cytoplasm | 0,001541 | 2,244981042 |
| P09429 | High mobility group protein B1 | Regulation of nucleotide metabolism | Nucleus | 0,019200 | 1,677061838 |
| P09471 | Guanine nucleotide-binding protein G(o) subunit alpha | Cell communication | Plasma membrane | 0,000006 | 2,175386959 |
| P09493 | Tropomyosin alpha-1 chain | Cell growth | Cytoskeleton | 0,006066 | 2,550099831 |
| P09622 | Dihydrolipoyl dehydrogenase_ mitochondrial | Metabolism; Energy pathways | Mitochondrion | 0,015963 | 1,753849584 |
| P09936 | Ubiquitin carboxyl-terminal hydrolase isozyme L1 | Protein metabolism | Cytoplasm | 0,000670 | 2,48447248 |
| P0DMV8 | Heat shock 70 kDa protein 1A | | Cytoplasm; Nucleus | 0,020307 | 1,439327653 |
| P0DP24 | Calmodulin-2 | Cell communication | Cytoplasm; Nucleus | 0,000143 | 11,36816528 |
| P10155 | 60 kDa SS-A/Ro ribonucleoprotein | Regulation of nucleotide metabolism | Cytoplasm | 0,011398 | 1,539351353 |
| P10599 | Thioredoxin | Metabolism; Energy pathways | Cytoplasm | 0,008225 | 1,925543807 |
| P10809 | 60 kDa heat shock protein_ mitochondrial | Protein folding | Mitochondrion | 0,000800 | 1,673308344 |
| P10909 | Clusterin | Immune response | Cytoplasm | 0,022458 | 2,180888369 |
| P11137 | Microtubule-associated protein 2 | Cell growth | Cytoplasm | 0,021320 | 1,562448768 |
| P11142 | Heat shock cognate 71 kDa protein | Protein metabolism | Cytoplasm | 0,000927 | 1,832670037 |
| P11216 | Glycogen phosphorylase_ brain form | Metabolism; Energy pathways | Cytoplasm | 0,003602 | 2,629010479 |
| P11532 | Dystrophin | Cell growth | Cytoplasm | 0,028536 | 1,627002621 |
| P11940 | Polyadenylate-binding protein 1 | Regulation of nucleotide metabolism | Cytoplasm; Nucleus | 0,002924 | 0,284578632 |
| P12268 | Inosine-5'-monophosphate dehydrogenase 2 | Metabolism; Energy pathways | Cytoplasm | 0,022160 | 1,311036617 |
| P12277 | Creatine kinase B-type | Metabolism; Energy pathways | Cytoplasm | 0,000945 | 2,567288508 |
| P12956 | X-ray repair cross-complementing protein 6 | Regulation of nucleotide metabolism | Nucleus | 0,047675 | 1,322064867 |
| P13667 | Protein disulfide-isomerase A4 | Protein metabolism | Endoplasmic reticulum | 0,009228 | 1,900324246 |
| P13861 | cAMP-dependent protein kinase type II-alpha regulatory subunit | Cell communication | Cytoplasm | 0,012335 | 1,849934911 |
| P14174 | Macrophage migration inhibitory factor | Cell communication | Extracellular | 0,000000 | 5,452247573 |
| P14314 | Glucosidase 2 subunit beta | Metabolism; Energy pathways | Endoplasmic reticulum | 0,039305 | 1,362863062 |
| P16152 | Carbonyl reductase [NADPH] 1 | Metabolism; Energy pathways | Cytoplasm | 0,041778 | 2,237760986 |
| P16401 | Histone H1.5 | Regulation of nucleotide metabolism | Nucleus | 0,046248 | 2,036540468 |
| P16949 | Stathmin | Cell growth | Cytoplasm | 0,000084 | 3,78954511 |
| P16989 | Y-box-binding protein 3 | Regulation of nucleotide metabolism | Nucleus | 0,022747 | 1,804122144 |
| P17844 | Probable ATP-dependent RNA helicase DDX5 | Regulation of nucleotide metabolism | Nucleus | 0,008272 | 1,449923646 |
| P17987 | T-complex protein 1 subunit alpha | Protein metabolism | Cytoplasm | 0,021912 | 2,466740118 |
| P21281 | V-type proton ATPase subunit B_ brain isoform | Transport | Endosome | 0,000010 | 1,558344576 |
| P22087 | rRNA 2'-O-methyltransferase fibrillarin | Regulation of nucleotide metabolism | Nucleus | 0,038911 | 0,519202443 |
| P22676 | Calretinin | Cell communication | Cytoplasm | 0,004405 | 4,810663523 |
| P23381 | Tryptophan--tRNA ligase_ cytoplasmic | Protein metabolism | Cytoplasm | 0,024157 | 0,623273137 |
| P23434 | Glycine cleavage system H protein_ mitochondrial | Metabolism; Energy pathways | Mitochondrion | 0,001400 | 1,911521005 |

| | | | | | |
|---|---|---|---|---|---|
| P23528 | Cofilin-1 | Cell growth | Cytoplasm | 0,014625 | 1,597391059 |
| P24539 | ATP synthase F(0) complex subunit B1_ mitochondrial | Metabolism; Energy pathways | Mitochondrion | 0,010465 | 0,540466061 |
| P25705 | ATP synthase subunit alpha_ mitochondrial | Metabolism; Energy pathways | Mitochondrion | 0,006083 | 2,845083522 |
| P25789 | Proteasome subunit alpha type-4 | Protein metabolism | Cytoplasm | 0,000010 | 2,103730426 |
| P26038 | Moesin | Cell growth | Cytoplasm | 0,000106 | 1,771643573 |
| P26368 | Splicing factor U2AF 65 kDa subunit | RNA metabolism | Cytoplasm; Nucleus | 0,032348 | 1,709346812 |
| P26378 | ELAV-like protein 4 | Regulation of nucleotide metabolism | Nucleus | 0,012280 | 4,418087716 |
| P26641 | Elongation factor 1-gamma | Protein metabolism | Cytoplasm | 0,003988 | 2,174778121 |
| P27695 | DNA-(apurinic or apyrimidinic site) lyase | Regulation of nucleotide metabolism | Cytoplasm | 0,000642 | 2,233503669 |
| P28070 | Proteasome subunit beta type-4 | Protein metabolism | Cytoplasm | 0,001951 | 1,817225795 |
| P28161 | Glutathione S-transferase Mu 2 | Metabolism; Energy pathways | Cytoplasm | 0,032607 | 0,122635942 |
| P30041 | Peroxiredoxin-6 | Metabolism; Energy pathways | Lysosome | 0,029738 | 1,465155069 |
| P30086 | Phosphatidylethanolamine-binding protein 1 | Cell communication | Cytoplasm | 0,000235 | 2,670603154 |
| P31323 | cAMP-dependent protein kinase type II-beta regulatory subunit | Cell communication | Cytoplasm | 0,032649 | 2,395304145 |
| P32119 | Peroxiredoxin-2 | Metabolism; Energy pathways | Cytoplasm | 0,000075 | 2,150091292 |
| P33176 | Kinesin-1 heavy chain | Cell growth | Mitochondrion | 0,027703 | 2,228651969 |
| P35232 | Prohibitin | Cell communication | Mitochondrion | 0,000221 | 1,569493465 |
| P35580 | Myosin-10 | Cell growth | Cytoplasm | 0,009957 | 1,640637533 |
| P35749 | Myosin-11 | Cell growth | Cytoplasm | 0,017217 | 1,551931028 |
| P35998 | 26S proteasome regulatory subunit 7 | Protein metabolism | Cytoplasm | 0,000417 | 2,19289918 |
| P36405 | ADP-ribosylation factor-like protein 3 | Cell communication | Cytoplasm | 0,000757 | 2,174710584 |
| P37108 | Signal recognition particle 14 kDa protein | Protein metabolism | Cytoplasm | 0,000015 | 1,732894835 |
| P37198 | Nuclear pore glycoprotein p62 | Transport | Nucleus | 0,000745 | 4,898039289 |
| P38159 | RNA-binding motif protein_ X chromosome | Regulation of nucleotide metabolism | Nucleus | 0,007735 | 3,010859592 |
| P39019 | 40S ribosomal protein S19 | Protein metabolism | Nucleus | 0,013595 | 1,74433533 |
| P39687 | Acidic leucine-rich nuclear phosphoprotein 32 family member A | Transcription | Endoplasmic reticulum | 0,006737 | 2,017895473 |
| P40227 | T-complex protein 1 subunit zeta | Protein metabolism | Cytoplasm | 0,039424 | 2,424869841 |
| P40925 | Malate dehydrogenase_ cytoplasmic | Metabolism; Energy pathways | Cytoplasm | 0,001429 | 2,682650974 |
| P40926 | Malate dehydrogenase_ mitochondrial | Metabolism; Energy pathways | Mitochondrion | 0,031708 | 1,352415375 |
| P42166 | Lamina-associated polypeptide 2_ isoform alpha | | | 0,014914 | 1,582368835 |
| P43487 | Ran-specific GTPase-activating protein | Cell communication | Cytoplasm | 0,034649 | 1,395639023 |
| P43686 | 26S proteasome regulatory subunit 6B | Protein metabolism | Nucleus | 0,000172 | 1,724158552 |
| P45974 | Ubiquitin carboxyl-terminal hydrolase 5 | Protein metabolism | Extracellular | 0,005219 | 0,5662373 |
| P46782 | 40S ribosomal protein S5 | Protein metabolism | Ribosome | 0,012027 | 1,438527599 |
| P46783 | 40S ribosomal protein S10 | Protein metabolism | Ribosome | 0,022989 | 0,279321508 |
| P46821 | Microtubule-associated protein 1B | Cell growth | Cytoplasm | 0,004604 | 5,87930572 |
| P48643 | T-complex protein 1 subunit epsilon | Protein metabolism | Centrosome | 0,008746 | 2,456875552 |
| P48723 | Heat shock 70 kDa protein 13 | Protein metabolism | Endoplasmic reticulum | 0,014553 | 1,586013019 |
| P49006 | MARCKS-related protein | Cell communication | | 0,031569 | 4,963617688 |
| P49327 | Fatty acid synthase | Metabolism; Energy pathways | Cytoplasm | 0,017385 | 1,627448611 |
| P49368 | T-complex protein 1 subunit gamma | Protein metabolism | Cytoplasm | 0,023073 | 1,664437048 |
| P49411 | Elongation factor Tu_ mitochondrial | Protein metabolism | Mitochondrion; Nucleus | 0,002477 | 1,608616627 |
| P49591 | Serine--tRNA ligase_ cytoplasmic | Metabolism; Energy pathways | Cytoplasm | 0,000170 | 6,085902228 |
| P49721 | Proteasome subunit beta type-2 | Protein metabolism | Cytoplasm | 0,023382 | 0,545905023 |
| P49915 | GMP synthase [glutamine-hydrolyzing] | Metabolism; Energy pathways | | 0,009031 | 0,458585862 |
| P50213 | Isocitrate dehydrogenase [NAD] subunit alpha_ mitochondrial | Metabolism; Energy pathways | Mitochondrion | 0,000494 | 0,643514233 |
| P50502 | Hsc70-interacting protein | Cell communication | Lysosome | 0,007255 | 2,548044191 |
| P50914 | 60S ribosomal protein L14 | Protein metabolism | Ribosome | 0,000237 | 1,678852154 |
| P51148 | Ras-related protein Rab-5C | Cell communication | Endosome | 0,020570 | 1,788211502 |
| P51784 | Ubiquitin carboxyl-terminal hydrolase 11 | Protein metabolism | Nucleus | 0,007860 | 2,627447501 |
| P51858 | Hepatoma-derived growth factor | Cell communication | Nucleus | 0,009932 | 1,966736821 |
| P52907 | F-actin-capping protein subunit alpha-1 | Cell growth | Cytoplasm | 0,023834 | 1,650442742 |
| P54136 | Arginine--tRNA ligase_ cytoplasmic | Protein metabolism | Cytoplasm | 0,001387 | 2,106776597 |
| P54577 | Tyrosine--tRNA ligase_ cytoplasmic | Metabolism; Energy pathways | Cytoplasm | 0,027478 | 1,422940258 |
| P54727 | UV excision repair protein RAD23 homolog B | Regulation of nucleotide metabolism | Cytoplasm; Nucleus | 0,019774 | 2,387665653 |
| P55072 | Transitional endoplasmic reticulum ATPase | Metabolism; Energy pathways | Cytoplasm | 0,016113 | 4,596795152 |
| P55209 | Nucleosome assembly protein 1-like 1 | Regulation of nucleotide metabolism | Cytoplasm; Mitochondrion | 0,041993 | 1,638856919 |
| P55854 | Small ubiquitin-related modifier 3 | Protein metabolism | Cytoplasm; Nucleus | 0,000017 | 2,902791705 |
| P56134 | ATP synthase subunit f_ mitochondrial | Metabolism; Energy pathways | Mitochondrion | 0,037842 | 0,385460168 |
| P59768 | Guanine nucleotide-binding protein G(I)/G(S)/G(O) subunit gamma-2 | Cell communication | | 0,000548 | 2,434908073 |
| P60174 | Triosephosphate isomerase | Metabolism; Energy pathways | Cytoplasm | 0,000083 | 3,913742323 |
| P60660 | Myosin light polypeptide 6 | Cell growth | Cytoplasm | 0,003606 | 2,591649563 |
| P60709 | Actin_ cytoplasmic 1 | Cell growth | Cytoplasm | 0,040687 | 2,932257282 |
| P61026 | Ras-related protein Rab-10 | Cell communication | Nucleus | 0,014351 | 1,750667202 |
| P61088 | Ubiquitin-conjugating enzyme E2 N | Protein metabolism | Nucleus | 0,000016 | 1,990869761 |
| P61266 | Syntaxin-1B | Transport | Plasma membrane | 0,028375 | 2,366590455 |
| P61586 | Transforming protein RhoA | Cell communication | Cytoplasm | 0,008164 | 2,983594687 |
| P61601 | Neurocalcin-delta | Cell communication | Cytoplasm | 0,030912 | 1,634865575 |
| P61604 | 10 kDa heat shock protein_ mitochondrial | Protein metabolism | Mitochondrion | 0,000009 | 2,778461817 |
| P61981 | 14-3-3 protein gamma | Cell communication | Cytoplasm | 0,048999 | 1,698868152 |
| P62191 | 26S proteasome regulatory subunit 4 | Protein metabolism | Cytoplasm | 0,000278 | 2,717973768 |
| P62269 | 40S ribosomal protein S18 | Protein metabolism | Ribosome | 0,024500 | 1,937082342 |
| P62304 | Small nuclear ribonucleoprotein E | Regulation of nucleotide metabolism | Nucleus | 0,004857 | 0,158825875 |
| P62328 | Thymosin beta-4 | Cell growth | Cytoplasm | 0,001050 | 5,688866399 |
| P62714 | Serine/threonine-protein phosphatase 2A catalytic subunit beta isoform | Cell communication | Cytoplasm; Nucleus | 0,032752 | 1,144638178 |
| P62829 | 60S ribosomal protein L23 | Protein metabolism | Ribosome | 0,017132 | 1,89542136 |
| P62917 | 60S ribosomal protein L8 | Protein metabolism | Nucleus | 0,007827 | 1,373061557 |
| P62937 | Peptidyl-prolyl cis-trans isomerase A | Protein folding | Cytoplasm; Mitochondrion | 0,000006 | 2,687671822 |
| P62987 | Ubiquitin-60S ribosomal protein L40 | Protein metabolism | Ribosome | 0,000040 | 2,659383552 |
| P63000 | Ras-related C3 botulinum toxin substrate 1 | Cell communication | Cytoplasm | 0,000248 | 1,676853813 |
| P63104 | 14-3-3 protein zeta/delta | Regulation of cell cycle | Cytoplasm | 0,002786 | 2,037317413 |
| P63208 | S-phase kinase-associated protein 1 | Protein metabolism | Nucleus | 0,018562 | 1,484630943 |
| P67809 | Nuclease-sensitive element-binding protein 1 | Regulation of nucleotide metabolism | Nucleus | 0,023294 | 1,868567001 |
| P67936 | Tropomyosin alpha-4 chain | Cell growth | Cytoskeleton | 0,014948 | 3,806361248 |
| P68036 | Ubiquitin-conjugating enzyme E2 L3 | Protein metabolism | Cytoplasm | 0,004306 | 4,940752027 |
| P68371 | Tubulin beta-4B chain | Cell growth | Cytoplasm | 0,043237 | 1,940299338 |
| P78559 | Microtubule-associated protein 1A | Cell growth | Cytoplasm | 0,000460 | 2,544974871 |
| P80404 | 4-aminobutyrate aminotransferase_ mitochondrial | Metabolism; Energy pathways | Mitochondrion | 0,027871 | 2,210104573 |
| P83916 | Chromobox protein homolog 1 | Regulation of nucleotide metabolism | Nucleus | 0,000477 | 1,967504985 |
| P84077 | ADP-ribosylation factor 1 | Signal transduction | Golgi apparatus | 0,029483 | 2,656101013 |
| P84103 | Serine/arginine-rich splicing factor 3 | Regulation of nucleotide metabolism | Nucleus | 0,004335 | 2,431538889 |
| Q00610 | Clathrin heavy chain 1 | Cell growth | Cytoplasm | 0,030669 | 1,908022019 |
| Q02790 | Peptidyl-prolyl cis-trans isomerase FKBP4 | Metabolism; Energy pathways | Cytoplasm | 0,001884 | 1,332204361 |

| | | | | | |
|---|---|---|---|---|---|
| Q04760 | Lactoylglutathione lyase | Metabolism; Energy pathways | Cytoplasm | 0,001383 | 2,092829595 |
| Q04917 | 14-3-3 protein eta | Cell communication | Cytoplasm | 0,049326 | 1,871833099 |
| Q05639 | Elongation factor 1-alpha 2 | Protein metabolism | Cytoplasm | 0,019878 | 1,455816713 |
| Q05682 | Caldesmon | Cell growth | Cytoplasm | 0,009880 | 2,705691034 |
| Q06323 | Proteasome activator complex subunit 1 | Protein metabolism | Nucleus | 0,004605 | 1,798822628 |
| Q06830 | Peroxiredoxin-1 | Metabolism; Energy pathways | Cytoplasm | 0,040068 | 1,941901288 |
| Q07021 | Complement component 1 Q subcomponent-binding protein_ mitochondrial | Immune response | Mitochondrion | 0,004835 | 1,517730954 |
| Q07866 | Kinesin light chain 1 | Signal transduction | Cytoplasm | 0,001460 | 1,729718449 |
| Q08043 | Alpha-actinin-3 | Cell growth | Cytoplasm | 0,000317 | 1,472062324 |
| Q09028 | Histone-binding protein RBBP4 | Regulation of nucleotide metabolism | Nucleus | 0,013135 | 1,320136313 |
| Q12765 | Secernin-1 | Immune response | Cytoplasm | 0,014923 | 3,078739534 |
| Q13098 | COP9 signalosome complex subunit 1 | Cell communication | Nucleus | 0,000101 | 1,94210178 |
| Q13153 | Serine/threonine-protein kinase PAK 1 | Cell communication | Plasma membrane | 0,004535 | 6,488271134 |
| Q13404 | Ubiquitin-conjugating enzyme E2 variant 1 | Cell differentiation | Nucleus | 0,007012 | 1,516709669 |
| Q13509 | Tubulin beta-3 chain | Cell growth | Cytoplasm | 0,009175 | 2,360523534 |
| Q13620 | Cullin-4B | Protein metabolism | Nucleus | 0,017783 | 3,704437199 |
| Q13885 | Tubulin beta-2A chain | | | 0,000246 | 1,965593106 |
| Q14008 | Cytoskeleton-associated protein 5 | Mitosis | Centrosome | 0,027123 | 1,50342356 |
| Q14011 | Cold-inducible RNA-binding protein | Cell communication | Nucleus | 0,009220 | 2,179301066 |
| Q14117 | Dihydropyrimidinase | Regulation of nucleotide metabolism | Cytoplasm | 0,000662 | 1,406999351 |
| Q14141 | Septin-6 | | Cytoplasm | 0,008088 | 2,934877417 |
| Q14195 | Dihydropyrimidinase-related protein 3 | Metabolism; Energy pathways | Cytoplasm | 0,000812 | 3,428140731 |
| Q14204 | Cytoplasmic dynein 1 heavy chain 1 | Metabolism; Energy pathways | Cytoplasm | 0,004551 | 1,44211235 |
| Q14240 | Eukaryotic initiation factor 4A-II | Protein metabolism | Cytoplasm | 0,000127 | 2,10519341 |
| Q14247 | Src substrate cortactin | Cell growth | Cytoplasm | 0,028210 | 2,004139481 |
| Q14257 | Reticulocalbin-2 | Cell communication | Endoplasmic reticulum | 0,029599 | 1,651999573 |
| Q14344 | Guanine nucleotide-binding protein subunit alpha-13 | Cell communication | Plasma membrane | 0,032823 | 1,149779435 |
| Q14498 | RNA-binding protein 39 | Regulation of nucleotide metabolism | Nucleus | 0,027667 | 1,522493816 |
| Q14576 | ELAV-like protein 3 | Regulation of nucleotide metabolism | Nucleus | 0,001719 | 0,09289286 |
| Q15058 | Kinesin-like protein KIF14 | Cell growth | Microtubule | 0,010381 | 0,162533191 |
| Q15063 | Periostin | Cell communication | | 0,003422 | 0,025271161 |
| Q15084 | Protein disulfide-isomerase A6 | Protein metabolism | Endoplasmic reticulum | 0,005007 | 2,420062454 |
| Q15102 | Platelet-activating factor acetylhydrolase IB subunit gamma | Metabolism; Energy pathways | Cytoplasm | 0,003112 | 1,759616448 |
| Q15185 | Prostaglandin E synthase 3 | Protein metabolism | Cytoplasm | 0,010390 | 1,704979434 |
| Q15293 | Reticulocalbin-1 | Cell communication | Endoplasmic reticulum | 0,000871 | 2,150265659 |
| Q15365 | Poly(rC)-binding protein 1 | Regulation of nucleotide metabolism | Nucleus | 0,048370 | 1,565494144 |
| Q15417 | Calponin-3 | Cell growth | Cytoplasm | 0,000044 | 2,574997371 |
| Q15435 | Protein phosphatase 1 regulatory subunit 7 | Regulation of nucleotide metabolism | Nucleus | 0,000767 | 5,935875578 |
| Q15631 | Translin | Regulation of nucleotide metabolism | Nucleus | 0,028200 | 1,437517269 |
| Q15700 | Disks large homolog 2 | Cell communication | Plasma membrane | 0,001751 | 1,772664327 |
| Q15843 | NEDD8 | Protein metabolism | Nucleus | 0,003854 | 4,456194812 |
| Q16181 | Septin-7 | Cell communication | Cytoskeleton | 0,019275 | 10,44709984 |
| Q16543 | Hsp90 co-chaperone Cdc37 | Protein metabolism | Cytoplasm | 0,000993 | 1,87419871 |
| Q16555 | Dihydropyrimidinase-related protein 2 | Cell communication | Cytoplasm | 0,000397 | 2,421927452 |
| Q16629 | Serine/arginine-rich splicing factor 7 | Regulation of nucleotide metabolism | Nucleus | 0,002531 | 2,250790013 |
| Q16658 | Fascin | Cell growth | Cytoplasm | 0,039815 | 1,644114619 |
| Q16698 | 2_4-dienoyl-CoA reductase_ mitochondrial | Metabolism; Energy pathways | Mitochondrion | 0,003434 | 0,55541008 |
| Q3ZCM7 | Tubulin beta-8 chain | Cell growth | | 0,017338 | 1,539927587 |
| Q53GQ0 | Very-long-chain 3-oxoacyl-CoA reductase | Metabolism; Energy pathways | Endoplasmic reticulum | 0,002213 | 0,652490858 |
| Q53GS9 | U4/U6.U5 tri-snRNP-associated protein 2 | Protein metabolism | Nucleus | 0,000088 | 1,59544169 |
| Q562R1 | Beta-actin-like protein 2 | | | 0,004534 | 1,453138889 |
| Q5JWF2 | Guanine nucleotide-binding protein G(s) subunit alpha isoforms XLas | | Cytoplasm | 0,030940 | 1,433181469 |
| Q6H8Q1 | Actin-binding LIM protein 2 | Cell growth | Cytoplasm | 0,002009 | 1,390526647 |
| Q6NXG1 | Epithelial splicing regulatory protein 1 | | Nucleus | 0,008787 | 0,271658095 |
| Q6P587 | Acylpyruvase FAHD1_ mitochondrial | | Mitochondrion | 0,014301 | 1,299265211 |
| Q6PCE3 | Glucose 1_6-bisphosphate synthase | Metabolism; Energy pathways | Cytoplasm | 0,005755 | 2,132171532 |
| Q6PEY2 | Tubulin alpha-3E chain | Cell growth | | 0,029850 | 1,631243281 |
| Q6PIW4 | Fidgetin-like protein 1 | Cell cycle | Nucleus | 0,004196 | 2,158070013 |
| Q6S8J3 | POTE ankyrin domain family member E | | Plasma membrane | 0,000036 | 5,261387464 |
| Q71UI9 | Histone H2A.V | Regulation of gene expression | Nucleus | 0,004956 | 2,007182756 |
| Q86Y82 | Syntaxin-12 | Transport | Endosome | 0,000452 | 1,871330097 |
| Q8IYB3 | Serine/arginine repetitive matrix protein 1 | Regulation of nucleotide metabolism | Nucleus | 0,016809 | 0,421057444 |
| Q8IYT4 | Katanin p60 ATPase-containing subunit A-like 2 | Metabolism; Energy pathways | | 0,042661 | 2,078808923 |
| Q8N0Y7 | Probable phosphoglycerate mutase 4 | Metabolism; Energy pathways | | 0,045956 | 0,571492472 |
| Q8N1G4 | Leucine-rich repeat-containing protein 47 | | Cytoplasm | 0,010122 | 1,458840997 |
| Q8N1N0 | C-type lectin domain family 4 member F | Cell communication | Plasma membrane | 0,000126 | 1,895356986 |
| Q8N283 | Ankyrin repeat domain-containing protein 35 | | | 0,012977 | 1,933920009 |
| Q8N446 | Zinc finger protein 843 | | | 0,024281 | 2,265420921 |
| Q8N4C6 | Ninein | Cell growth | Centrosome | 0,003816 | 1,458044438 |
| Q8N568 | Serine/threonine-protein kinase DCLK2 | Cell communication | | 0,041562 | 2,306858292 |
| Q8N5K1 | CDGSH iron-sulfur domain-containing protein 2 | | Plasma membrane | 0,000148 | 2,480130777 |
| Q8N6D5 | Ankyrin repeat domain-containing protein 29 | | | 0,001532 | 2,279329882 |
| Q8NB46 | Serine/threonine-protein phosphatase 6 regulatory ankyrin repeat subunit C | | | 0,004422 | 9,057306503 |
| Q8NFI4 | Putative protein FAM10A5 | | | 0,008676 | 2,136001501 |
| Q8WXF1 | Paraspeckle component 1 | Regulation of nucleotide metabolism | Nucleus | 0,006365 | 1,855020049 |
| Q8WZA9 | Immunity-related GTPase family Q protein | | | 0,000798 | 3,349862763 |
| Q92688 | Acidic leucine-rich nuclear phosphoprotein 32 family member B | | Nucleus | 0,001681 | 1,826826651 |
| Q92734 | Protein TFG | | | 0,000041 | 1,623976204 |
| Q92804 | TATA-binding protein-associated factor 2N | Signal transduction | Cytoplasm | 0,014965 | 4,179453218 |
| Q92841 | Probable ATP-dependent RNA helicase DDX17 | Regulation of nucleotide metabolism | Nucleus | 0,033428 | 1,583202872 |
| Q92878 | DNA repair protein RAD50 | Regulation of nucleotide metabolism | Nucleus | 0,000316 | 2,789032089 |
| Q93045 | Stathmin-2 | Cell communication | Golgi apparatus | 0,001125 | 3,36085983 |
| Q96JE9 | Microtubule-associated protein 6 | Cell growth | Cytoskeleton | 0,007744 | 3,79347076 |
| Q96M83 | Coiled-coil domain-containing protein 7 | | | 0,007578 | 1,667058536 |
| Q96P16 | Regulation of nuclear pre-mRNA domain-containing protein 1A | Regulation of nucleotide metabolism | Nucleus | 0,017844 | 0,491072812 |
| Q99497 | Protein/nucleic acid deglycase DJ-1 | Regulation of nucleotide metabolism | Cytoplasm | 0,031686 | 2,840061794 |
| Q99536 | Synaptic vesicle membrane protein VAT-1 homolog | Transport | Cytoplasm | 0,049415 | 2,294719181 |
| Q99733 | Nucleosome assembly protein 1-like 4 | Regulation of nucleotide metabolism | Nucleus | 0,000205 | 1,726657824 |
| Q99961 | Endophilin-A2 | Cell communication | Cytoplasm; Nucleus | 0,021414 | 1,30328806 |
| Q9BPW8 | Protein NipSnap homolog 1 | | Mitochondrion | 0,031444 | 1,304884688 |
| Q9BS26 | Endoplasmic reticulum resident protein 44 | Protein metabolism | Endoplasmic reticulum | 0,010718 | 1,257953912 |
| Q9BSA4 | Protein tweety homolog 2 | | Plasma membrane | 0,007477 | 1,44302528 |
| Q9BUF5 | Tubulin beta-6 chain | Cell growth | | 0,008748 | 2,082584317 |

| | | | | | |
|---|---|---|---|---|---|
| Q9BUT1 | 3-hydroxybutyrate dehydrogenase type 2 | Metabolism; Energy pathways | | 0,039248 | 1,759711645 |
| Q9BVA1 | Tubulin beta-2B chain | Cell growth | | 0,001906 | 1,853167562 |
| Q9BYT8 | Neurolysin_ mitochondrial | Protein metabolism | Cytoplasm | 0,012099 | 2,045797909 |
| Q9C040 | Tripartite motif-containing protein 2 | | Cytoplasm | 0,000328 | 3,518496209 |
| Q9H0C8 | Integrin-linked kinase-associated serine/threonine phosphatase 2C | Cell death | Cytoplasm | 0,049368 | 0,279169919 |
| Q9H1E3 | Nuclear ubiquitous casein and cyclin-dependent kinase substrate 1 | Regulation of nucleotide metabolism | Nucleus | 0,000901 | 2,797512718 |
| Q9H9B4 | Sideroflexin-1 | Transport | Mitochondrion | 0,004657 | 0,772798774 |
| Q9HB07 | UPF0160 protein MYG1_ mitochondrial | | Cytoplasm | 0,001033 | 1,315226246 |
| Q9HC38 | Glyoxalase domain-containing protein 4 | | Mitochondrion | 0,023117 | 2,22854275 |
| Q9NQP4 | Prefoldin subunit 4 | Protein metabolism | Cytoplasm | 0,018024 | 2,413012791 |
| Q9NR31 | GTP-binding protein SAR1a | Cell communication | Endoplasmic reticulum | 0,001929 | 0,042692402 |
| Q9NRR5 | Ubiquilin-4 | Protein metabolism | Nucleus | 0,019276 | 0,512314356 |
| Q9NUD5 | Zinc finger CCHC domain-containing protein 3 | Regulation of nucleotide metabolism | | 0,007861 | 0,336560004 |
| Q9NVA2 | Septin-11 | Cell cycle | Cytoplasm | 0,000693 | 4,199053245 |
| Q9NYF8 | Bcl-2-associated transcription factor 1 | Regulation of nucleotide metabolism | Nucleus | 0,047403 | 0,226310547 |
| Q9NZI8 | Insulin-like growth factor 2 mRNA-binding protein 1 | Regulation of nucleotide metabolism | Cytoplasm | 0,044945 | 1,601448506 |
| Q9P258 | Protein RCC2 | Cell growth | Nucleus | 0,021411 | 0,15633396 |
| Q9P2K5 | Myelin expression factor 2 | Regulation of nucleotide metabolism | Nucleus | 0,024166 | 1,772039706 |
| Q9P2M7 | Cingulin | Cell growth | Nucleus | 0,000639 | 2,652951999 |
| Q9UEE9 | Craniofacial development protein 1 | Cell growth | Cytoplasm | 0,000366 | 5,036446775 |
| Q9UHD8 | Septin-9 | Cell proliferation | Cytoskeleton | 0,008033 | 0,172771629 |
| Q9UHD9 | Ubiquilin-2 | Protein metabolism | Cytoplasm | 0,002383 | 3,152110424 |
| Q9UHV9 | Prefoldin subunit 2 | Protein metabolism | | 0,000001 | 3,845595515 |
| Q9UI15 | Transgelin-3 | | Cytoplasm | 0,000331 | 3,063017519 |
| Q9UK76 | Jupiter microtubule associated homolog 1 | Cell communication | | 0,001224 | 2,367420024 |
| Q9UMX0 | Ubiquilin-1 | Protein metabolism | Cytoplasm | 0,005438 | 2,957739768 |
| Q9UNZ2 | NSFL1 cofactor p47 | Cell growth | Nucleus | 0,003702 | 1,944052929 |
| Q9Y230 | RuvB-like 2 | Regulation of nucleotide metabolism | Nucleus | 0,002725 | 1,329116946 |
| Q9Y265 | RuvB-like 1 | Regulation of nucleotide metabolism | Nucleus | 0,042735 | 1,604672961 |
| Q9Y277 | Voltage-dependent anion-selective channel protein 3 | Transport | Mitochondrion | 0,040515 | 2,303247957 |
| Q9Y281 | Cofilin-2 | Cytoskeleton organization | Cytoplasm | 0,000202 | 1,691703183 |
| Q9Y3E1 | Hepatoma-derived growth factor-related protein 3 | Cell growth | Nucleus | 0,010543 | 1,494147481 |
| Q9Y3E1 | Peptidyl-prolyl cis-trans isomerase A-like 4A | | | 0,000958 | 2,183576372 |
